# Developmental resilience of synaptome architecture

**DOI:** 10.1101/2021.12.21.473638

**Authors:** Laura Tomas-Roca, Zhen Qiu, Erik Fransén, Ragini Gokhale, Edita Bulovaite, David J. Price, Noboru H. Komiyama, Seth G.N. Grant

## Abstract

Neurodevelopmental disorders of genetic origin delay the acquisition of normal abilities and cause disabling phenotypes. Spontaneous attenuation and even complete amelioration of symptoms in early childhood and adolescence occur in many disorders^1–10^, suggesting that brain circuits possess an intrinsic capacity to repair themselves. We examined the molecular composition of almost a trillion excitatory synapses on a brain-wide scale between birth and adulthood in mice carrying a mutation in the homeobox transcription factor *Pax6*, a neurodevelopmental disorder model^11^. *Pax6* haploinsufficiency had no impact on total synapse number at any age. By contrast, the postnatal expansion of synapse diversity and acquisition of normal synaptome architecture were delayed in all brain regions, interfering with network and cognitive functions. Specific excitatory synapse types and subtypes were affected in two key developmental age-windows. These phenotypes were reversed within 2-3 weeks of onset, restoring synaptome architecture to its normal developmental trajectory. Synapse subtypes with high rates of protein turnover mediated these events. These results show synaptome remodelling confers resilience to neurodevelopmental disorders.

---

Excitatory synapses, which make up the vast majority of synapses in the brain, have highly diverse identities resulting from their differing protein composition and protein lifetimes^12–14^. The anatomical distribution of these molecularly-diverse synapses can be studied using synaptome mapping technology, a large-scale, single-synapse resolution, systematic image analysis approach that has uncovered brain-wide excitatory synapse diversity in the mouse^12–14^. The Mouse Lifespan Synaptome Atlas^13^ revealed that synapse composition of the brain changes continuously across the lifespan, with trajectories of excitatory synapse types and subtypes in dendrites, neurons, circuits and brain regions, together defining the Lifespan Synaptome Architecture (LSA)^13^. A key feature of the LSA is its three distinct age-windows (LSA I-III), which correspond to childhood, adolescence/young adulthood, and the ageing adult.

Although the genetic mechanisms controlling synaptome architecture are only beginning to be understood, analysis of the spatial patterning of synapses suggests that developmental mechanisms controlling the body plan might be involved^14^. Transcription factors containing homeodomains play a key role in establishing the body plan and development of the brain^15–21^ and we hypothesized that they may control synaptome architecture. Pax6, a member of the homeodomain transcription factor family, is known to regulate the expression of PSD95 and SAP102^*22*^, which are postsynaptic proteins in excitatory synapses used in synaptome mapping^13,14^. Heterozygous (haploinsufficient) loss-of-function *PAX6* mutations cause autism, intellectual disability, epilepsy and aniridia (WAGR syndrome)^11,23–29^, which are phenocopied in *Pax6*^+/−^ mice^11,30,31^. Pax6 is expressed during embryogenesis in progenitor cells giving rise to forebrain, midbrain and hindbrain structures, and in postnatal ages in subsets of diencephalic neurons^32,33^. Our investigations into the role of Pax6 not only show that development of the synaptome architecture of the brain is organized by homeobox genes, but also that the synaptome has a remarkable capacity to repair itself during childhood and adolescence.

## Mapping the developing synaptome architecture of *Pax6*^+/−^ mice

The synaptome architecture of *Pax6*^+/−^ mice was analyzed using the SYNMAP synaptome mapping pipeline^13,14^, which quantifies the expression of postsynaptic proteins PSD95 and SAP102 in synaptic puncta (Fig S1). The levels of PSD95 and SAP102 are functionally important because they physically assemble proteins controlling synaptic transmission, plasticity and neuronal excitability into macromolecular complexes^34–37^ and altering their expression leads to changes in synaptic and cognitive functions^38^,^39^. To genetically label PSD95 and SAP102 in excitatory synapses, *Pax6*^+/−^ mice were crossed with PSD95-eGFP and SAP102-mKO2 mice to generate cohorts of *Pax6*^+/−^;*Psd95*^eGFP/eGFP^;*Sap102*^mKO2/mKO2^ and control (*Pax6*^+/+^;*Psd95*^eGFP/eGFP^;*Sap102*^mKO2/mKO2^) mice. Parasagittal brain sections were collected at day one (P1) and at one (P7), two (P14), three (P21), four (P28), five (P35), six (P42), seven (P49) and eight (P56) weeks. The first five time points correspond to LSA-I (P1-P28) and the latter four to LSA-II (P35-P56)^13^. Brain sections were imaged at single-synapse resolution on a spinning disk confocal microscope (pixel size 84 x 84 nm and optical resolution ~260 nm) and the density, intensity, size and shape parameters of individual puncta were acquired in 131 brain subregions. We classified the synaptic puncta into three types (type 1 express PSD95 only, type 2 express SAP102 only, and type 3 express both PSD95 and SAP102) and a further 37 subtypes on the basis of molecular and morphological features^14^. All data were registered to the Allen Developing Mouse Brain Atlas (Table S1) and are available (Table S2-S15) at Edinburgh DataShare^40^. We created the Pax6 Developmental Synaptome Atlas (https://brain-synaptome.org/Pax6_Developmental_Synaptome_Atlas/)^41^, an interactive visualization tool for displaying the spatial framework of datasets and differences between control and *Pax6*^+/−^ mice.

## Transient synaptome phenotypes in two age-windows

We measured a total of 3.65 x 10^11^ excitatory synapses and found no significant differences in synapse number between *Pax6*^+/−^ and control mice in any brain region or in the whole brain at any age (P > 0.05, Benjamini-Hochberg corrected). To test whether the LSA was impacted, we examined the density of the three synapse types and 37 subtypes at the nine ages (Fig 1, S2). At birth (P1) the *Pax6*^+/−^ synaptome was largely normal, but during the second and third postnatal weeks (P7-P21) strong synaptome phenotypes emerged in most brain regions, which then reverted to normal in week four (P28). Synaptome phenotypes then remerged in week five (P35-P42) before reverting again to normal by P49. This temporal progression of synaptome phenotypes describes two ‘phenotype waves’, one starting in the second postnatal week and the other in the sixth postnatal week, each lasting approximately 2 weeks. At the peak of each phenotype wave, synapse composition was affected in almost every region and subregion of the *Pax6*^+/−^ brain (Fig 1, S2).

**Figure 1.**
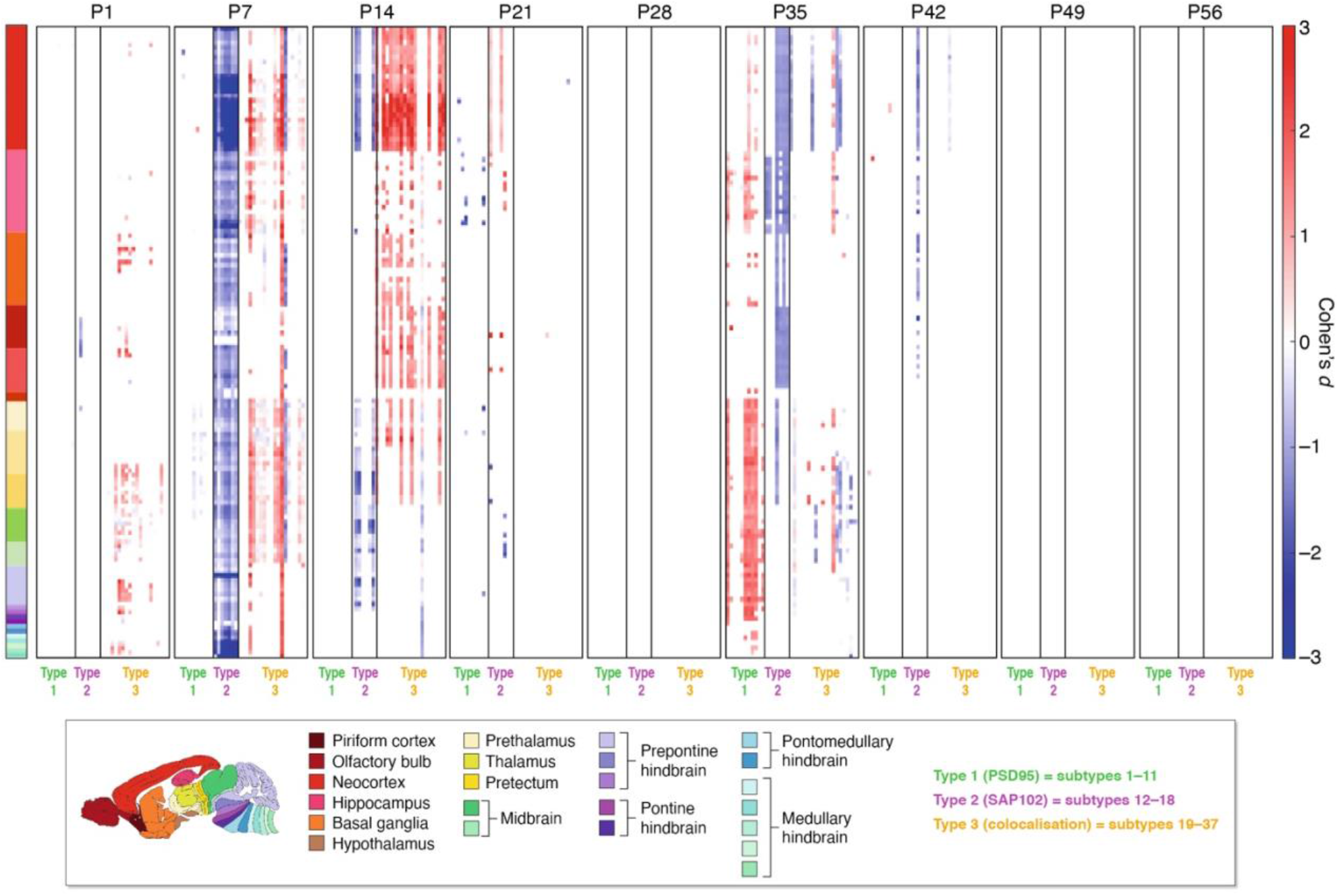
Synaptome phenotypes in *Pax6*^+/−^ mice. The difference (Cohen’s *d*) in synapse type and subtype density in 131 brain subregions at nine ages from birth to P56 between *Pax6*^+/−^ and control mice. Significantly different (P < 0.05, Benjamini-Hochberg corrected) subregions are shown. High-resolution graphs are provided in Fig S2.

## *Pax6* mutations disrupt brain network structure and function

The synaptome architecture of the brain can be described as a network of brain regions^14^, which has previously been shown to correlate with structural connectome networks and with dynamic network activity measured using resting-state fMRI^14^. To assess the impact of the *Pax6*^+/−^ mutation on brain synaptome networks, we examined the similarity of the synaptome between all brain regions at each age and the topology of networks built from the similarity matrices (Fig 2, S3). The similarity matrices of control mice at birth show high and homogeneous similarity, followed by a rapid decline by P14, consistent with previous results^13^. By contrast, in *Pax6*^+/−^ mice this decline was delayed, resulting in major differences of similarity matrix and, therefore, in the similarity ratio compared with control mice at P14 (Cohen’s *d* = 4.27 and P<0.005, Bayesian test with Benjamini-Hochberg correction, Fig 2A, B, S3). This developmental delay was overcome in the following week (P>0.05, Bayesian test with Benjamini-Hochberg correction, Fig 2A, B, S3), corresponding to the end of LSA-I. Two weeks later, at P35 (in LSA-II), there was another significant increase in the similarity of brain regions in *Pax6*^+/−^ mice (Cohen’s *d* = 1.24 and P<0.05 in similarity ratio, Bayesian test with Benjamini-Hochberg correction, Fig 2A, B, S3), which again was followed by a return to control values in the subsequent two weeks. When we examined the topology of synaptome networks using an index of small worldness^13,14^, this showed major reductions in small worldness at P14 (Cohen’s *d* = - 3.14 and P<0.05, Bayesian test with Benjamini-Hochberg correction, Fig 2C) and P35 (Cohen’s *d* = −4.48 and P<0.05, Bayesian test with Benjamini-Hochberg correction, Fig 2C) in the *Pax6*^+/−^ brain. These results indicate that the network properties of the brain are transiently impaired during the two phenotype waves.

**Figure 2.**
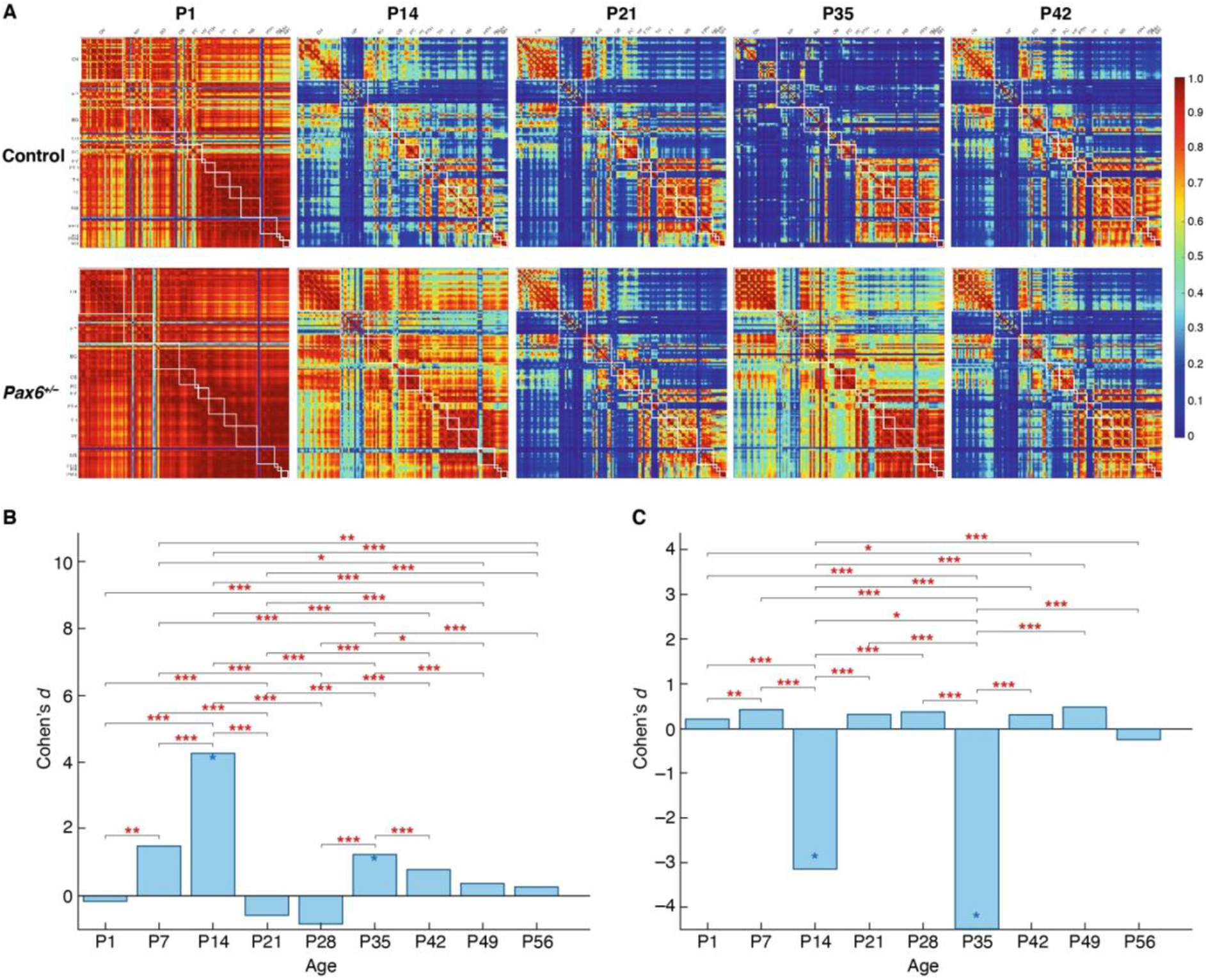
Synaptome architecture is disrupted in *Pax6*^+/−^ mice. A. Matrix heatmap of similarities between pairs of brain subregions (rows and columns) at the indicated ages in control (top row) and *Pax6*^+/−^ (bottom row) mice (see Fig S3 for all ages). White boxes indicate the subregions that belong to the same main brain region (see region list in Fig 1). Scale bar, similarity values. B. Differences (Cohen’s *d*) in similarity ratio between *Pax6*^+/−^ and control mice for the different age groups. A significant increase of ratio in *Pax6*^+/−^ mice is shown at P14 and P35: blue asterisk P<0.05, Bayesian test with Benjamini-Hochberg correction. The difference in ratio at P14 is significantly larger than at other ages: red asterisks, * P< 0.05, **P < 0.01, ***P < 0.001, two-way ANOVA with post-hoc multiple comparison test. C. Differences (Cohen’s *d*) in average small worldness between *Pax6*^+/−^ and control mice for the different age groups. A significant decrease of small worldness in *Pax6*^+/−^ mice is shown at P14 and P35: blue asterisk P<0.05, Bayesian test with Benjamini-Hochberg correction. The difference in small worldness at P14 is significantly larger than at other ages: red asterisks, * P< 0.05, **P < 0.01, ***P < 0.001, two-way ANOVA with post-hoc multiple comparison test.

To explore the functional consequences of synaptome phenotypes for synaptic electrophysiological properties relevant to the storage and recall of behavioral representations, we employed a computational simulation approach^13,14^ in which the synaptome of CA1 pyramidal neurons is stimulated with patterns of neural activity and the spatial output of excitatory postsynaptic potentials (EPSP) is quantified at P1, P7, P28, P35 and P56 (Fig 3A, S4, Table S14, S15). Theta burst and gamma train stimulation (Fig 3B), but not theta train or gamma burst (Fig S4), resulted in significant phenotypes at P7 and P35 in *Pax6*^+/−^ mice (P < 0.01, paired t-test, Benjamini-Hochberg corrected). These findings indicate that the synaptome in the CA1 region, which is a crucial structure for spatial navigation, learning and memory^42^, is reversibly impaired during the two phenotype waves in *Pax6*^+/−^ mice.

**Figure 3.**
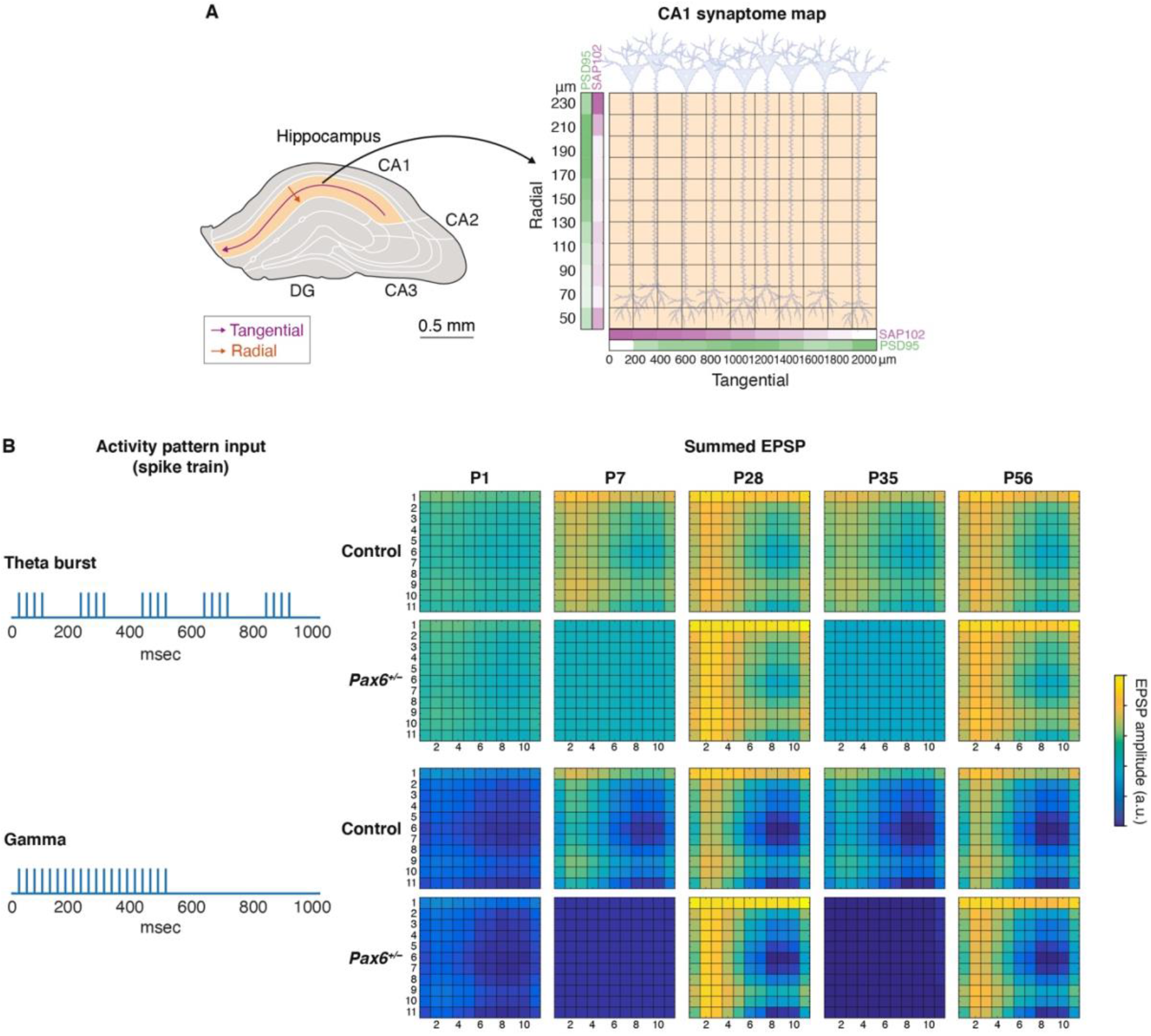
Synaptic responses to physiological spike patterns. A. Synaptic gradients along the radial and tangential directions, with gradient intensity symbolized by color intensity; PSD95 (green) and SAP102 (magenta) were used as in previous work^13,14^ to set synaptic properties of the computational model. Synaptic amplitudes of age groups (P1-P56) and genotype (control, *Pax6*^+/−^) were scaled based on intensity (Table S14, S15). B. Synapses were activated by spike patterns representing theta burst (top) and gamma frequency (bottom) activity. Summed EPSP response amplitudes were quantified (color bar, arbitrary units) and statistical differences between synaptic responses of control (upper) and *Pax6*^+/−^ (lower) were assessed.

## Synapses with rapid protein turnover mediate synaptome changes

To understand the mechanisms underpinning the dysfunction in brain networks during the phenotype waves and how this dysfunction is reversed, we examined the impact of the *Pax6*^+/−^ mutation on excitatory synapse type and subtype diversity. As shown in Figure 1, synapse types and subtypes were differentially affected during the phenotype waves. In both waves there was a loss of type 2 synapses (which express SAP102 only) and an increase in type 1 and 3 synapses (which express PSD95), suggesting that the PSD95-expressing synapses are compensating or adapting to the loss of SAP102-expressing synapses. Among the 30 PSD95-expressing synapse subtypes in our excitatory synapse catalog^14^, some were widely impacted throughout the *Pax6*^+/−^ brain, whereas others were affected in very few brain regions (Fig 4A). To further investigate how these changing numbers of synapse subtypes affect the synapse composition of each brain subregion at different ages, we asked if synapse diversity^13,14^ was altered. As shown in Figure 4C, there was a reduction in synapse diversity across most brain regions during the first phenotype wave (34/131 subregions in P1, 117/131 in P7 and 63/131 in P14), with little effects at any later ages.

**Figure 4.**
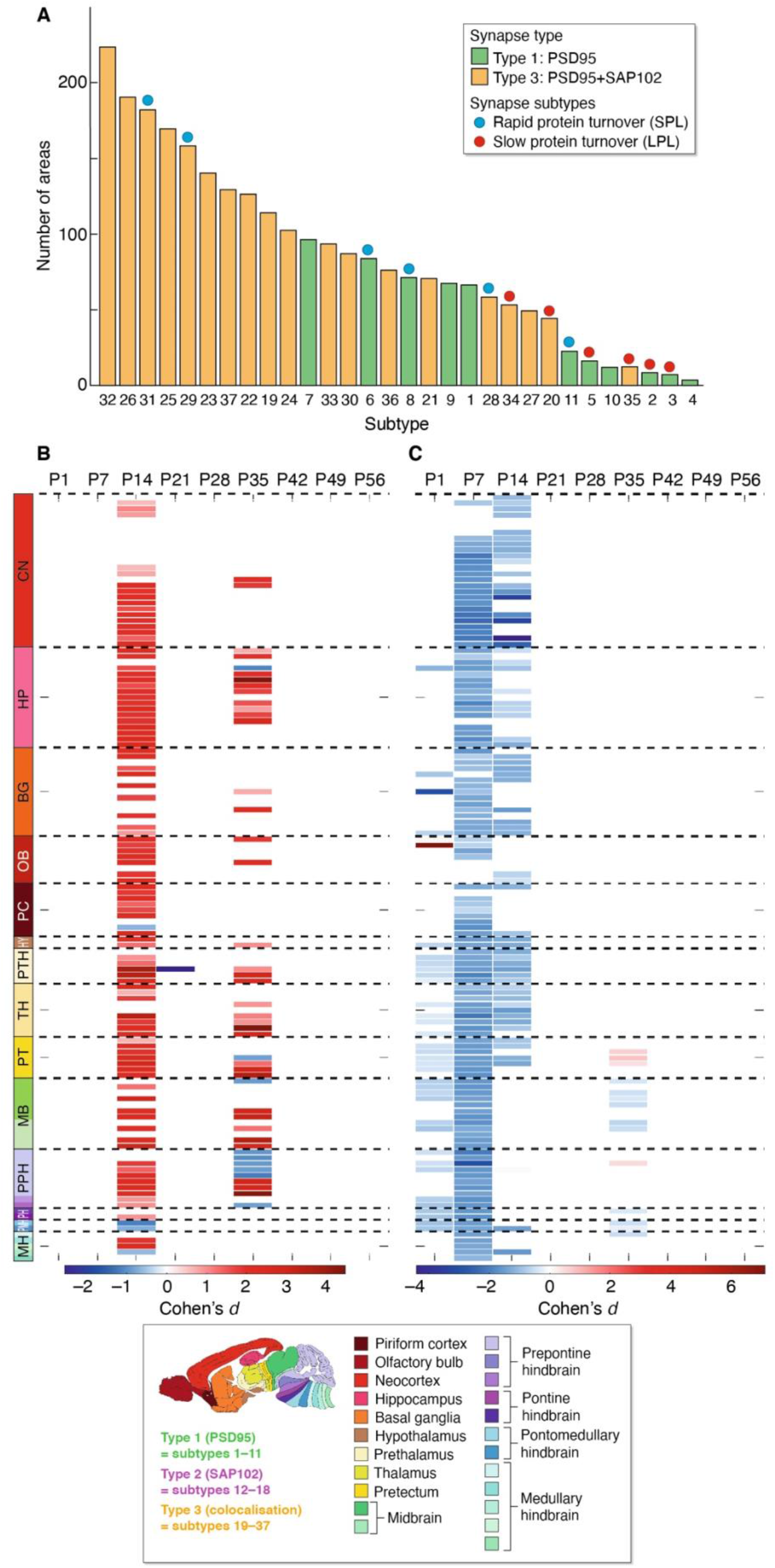
Synapse subtypes with rapid protein turnover are affected in *Pax6*^+/−^ mice. A. Ranking of affected type 1 and 3 synapse subtypes in *Pax6*^+/−^ mice. For each subtype the number of brain subregions that showed a significant difference (P< 0.05, Bayesian test with Benjamini-Hochberg correction) in density between control and *Pax6*^+/−^ mice was counted at all ages. Note skewing in SPL and LPL subtypes. B. Significantly larger phenotypes were observed in SPL synapses compared with LPL synapses in *Pax6*^+/−^ mice (Cohen’s *d*, scale bar indicates larger mutant phenotype of SPL types in red and smaller mutant effect of SPL types in blue; P < 0.05, Bayesian method with Benjamini-Hochberg correction) at P14 and P35. C. Changes in synapse diversity in brain subregions between *Pax6*^+/−^ and control mice for the different age groups. Heatmap shows the significant decrease (Cohen’s *d*, scale bar indicates increase in red and decrease in blue; P < 0.05, Bayesian method with Benjamini-Hochberg correction) of synapse diversity in *Pax6*^+/−^ mice in many brain subregions at P1, P7 and P14. Absence of color indicates subregions where no significant differences occurred.

We recently found that PSD95-expressing synapse subtypes can differ greatly in their rate of protein turnover, and classified them as short protein lifetime (SPL) (subtypes 6, 8, 11, 29, 28, 31) and long protein lifetime (LPL) (subtypes 2, 34, 3, 35, 5, 20) synapses^12^. Because protein turnover is important for maintenance and remodeling of the proteome we asked if SPL and LPL synapses are differentially involved with the dynamic synaptome phenotypes in *Pax6*^+/−^ mice. We found that SPL synapses are disproportionately affected in *Pax6*^+/−^ mice, showing a significantly larger phenotype than LPL synapses (P< 0.05, Bayesian test with Benjamini-Hochberg correction) in 72% (94/131) of subregions at P14 and 28% (36/131) of subregions at P35 (Fig 4B). By contrast, only 3% (4/131) of subregions at P14 and 7% (9/131) of subregions at P35 showed a significant larger phenotype in LPL over SPL synapses (P< 0.05, Bayesian test with Benjamini-Hochberg correction) (Fig 4B). These results show the dynamic and transient nature of these phenotypes resides largely with SPL synapses, whereas LPL synapses remain more resilient to the impact of this genetic perturbation.

## Discussion

We have assessed the effects of a germline mutation in *Pax6*, a homeobox transcription factor, on the development of the synaptome architecture of the brain and created the Pax6 Developmental Synaptome Atlas^41^. We analyzed almost a trillion individual excitatory synapses at weekly intervals from birth to maturity at 2 months of age and found that the total number of excitatory synapses was not affected in any brain region at any age in *Pax6*^+/−^ mice. Instead, there are striking changes in the regional composition of excitatory synapse types and subtypes. Remarkably, these synaptome phenotypes emerged transiently in two age-windows, indicating that the molecular identity of excitatory synapses is dynamic and capable of restoring the normal developmental trajectory of the LSA in a genetic neurodevelopmental disorder.

The restoration of the LSA required changes in the numbers of synapse types and subtypes, indicating that their postsynaptic proteomes were being remodeled. Protein turnover is a fundamental mechanism for homeostatic maintenance of the proteome (proteostasis) and is required for proteome remodeling instructed by transcriptional programs^43–45^. Consistent with this, in *Pax6*^+/−^ mice the density of synapse subtypes with the fastest protein turnover (SPL synapses) were most affected during the age-windows. Moreover, the protein turnover rate in SPL synapses^12^ corresponds well with the duration of the synaptome repair in *Pax6*^+/−^ mice.

The age-windows during which the LSA was restored in *Pax6*^+/−^ mice correspond to two important transitions in the life of a mouse: from dependency on the mother to independent feeding in LSA-I, and the attainment of sexual maturity and adult behaviors in LSA-II. By ensuring normal trajectories of synaptome architecture during these crucial transitions, maladaptive behaviors caused by underlying genetic variation would be minimized and the mice more likely to survive. The brain of young animals is highly enriched in SPL synapses^12^, providing a pool of synapse subtypes capable of rapidly remodeling and repairing the LSA during development.

In humans, neurodevelopmental disorders delay the acquisition of speech and language, social interactions, learning, attention and motor skills, and also manifest with the onset of epilepsy or motoric dysfunction. Despite this, spontaneous attenuation and even complete amelioration of symptoms occur in early childhood and adolescence in some individuals^1–10^. This amelioration affects behaviors controlled by different brain regions arising from diverse kinds of mutations, indicating that the capacity for spontaneous recovery from neurodevelopmental disorders is a pervasive and general mechanism in the developing nervous system. This capacity in humans and the findings of the present study are reminiscent of the concept of canalization introduced by Waddington almost 80 years ago^46^. Canalization provides resilience and maintains the normal trajectories of developmental programs in the face of genetic or environmental perturbations and has been invoked as an explanation for why apparently normal individuals carry deleterious mutations, why symptom penetrance varies in diseases^47–53^, and how developing neuronal networks overcome mutations^53^. We suggest that synapse diversity and rapidly remodeling synapses provide a capacity for canalization in the brain. The dynamic and flexible nature of excitatory synapse diversity and synaptome architecture suggests that therapeutic approaches, potentially targeting SPL synapse subtypes, might enhance resilience to neurodevelopmental disorders and environmental insults.

## Acknowledgments

**General**

C. McLaughlin, E. Sigfridsson, B. Notman, R. Dahan, G. Varga, R. Gokhale, B. Koniaris, A. Bujalance and N. Skene for advice and technical assistance. C. Davey for editing. D. Maizels for artwork.

## Author contributions

LTR planned the experiments, collected, imaged and analyzed brain samples, performed image and data analyses. ZQ developed software and performed image and data analyses. EF analyzed data and performed computational modeling of neural activity. EB provided data on synapse protein turnover. NHK and DJP provided supervision. SGNG supervised the project and wrote the manuscript.

## Data and materials availability

Data are available at Edinburgh DataShare^40^.

