## Supplementary Material for "Developmental resilience of synaptome architecture"

### Contents:

Material and Methods

Supplementary Figures (S1-4)

Supplementary Table legends (S1-15)

### Materials and Methods

#### Animals

Animal procedures were performed in accordance with UK Home Office regulations and approved by the Edinburgh University Director Biological Services. Generation and characterization of *Psd95*$^{eGFP/eGFP}$;*Sap102*$^{mKO2/mKO2}$ knock-in mice was described previously[14]. *Pax6*$^{+/-}$ mice[54] were crossed with PSD95-eGFP and SAP102-mKO2 mice to generate cohorts of *Pax6*$^{+/-}$;*Psd95*$^{eGFP/eGFP}$;*Sap102*$^{mKO2/mKO2}$ and control mice (*Pax6*$^{+/+}$;*Psd95*$^{eGFP/eGFP}$;*Sap102*$^{mKO2/mKO2}$). Both control (c) and mutant (m) mice from both sexes were collected at nine postnatal time points: one (P1, c=11, m=6), seven (P7, c=7, m=7), fourteen (P14, c=6, m=7), twenty-one (P21, c=7, m=8), twenty-eight (P28, c=8, m=7), thirty-five (P35, c=11, m=6), forty-two (P42, c=7, m=6), forty-nine (P49, c=6, m=6) and fifty-six (P56, c=16, m=9) days.

#### Tissue collection and sectioning

Mice were anesthetized by an intraperitoneal injection of 20% (w/v) sodium pentobarbital (Euthatal, Merial Animal Health or Pentoject, Animalcare: 0.01 ml for P1-P7, 0.05 ml for P14-P21, 0.1 ml for P28-P56). After complete anesthesia, phosphate-buffered saline (PBS, Oxoid; 5 ml for P1-P14, 10 ml for P21-P56) was perfused transcardially, followed by 4% (v/v) paraformaldehyde (PFA, Alfa Aesar; 5 ml for P1-P14, 10 ml for P21-P56). Whole brains were dissected out and immediately postfixed at 4°C in 4% PFA (2 h P1-P14, 4 h P21-P56) before transferring them into 30% (w/v) sucrose at 4°C (in 1×PBS, VWR Chemicals). Brains were then embedded into Optimal Cutting Temperature (OCT,

CellPath) medium within a cryomould and frozen in isopentane cooled with liquid nitrogen. Brains were then sectioned in the parasagittal plane at 18 μm thickness using an NX70 cryostat (Thermo Fisher Scientific). Cryosections were mounted on Superfrost Plus glass slides (Thermo Fisher Scientific) and stored at −80°C.

**Tissue preparation**

Parasagittal sections from left hemisphere (corresponding to sections 12-13/24 from sagittal Allen Brain Reference Atlas)[55] were washed for 5 min in PBS, incubated for 15 min in 1 mg/ml DAPI (Sigma), washed with PBS and mounted using home-made MOWIOL (Calbiochem) containing 2.5% anti-fading agent DABCO (Sigma-Aldrich), covered with a coverslip (thickness #1.5, VWR International) and imaged the following day.

**Spinning disk confocal microscopy**

Fast high-resolution imaging was achieved using an Andor Revolution XDi system equipped with an Olympus UPlanSAPO 100x oil-immersion lens (NA 1.4), a CSU-X1 spinning-disk (Yokogawa) and an Andor iXon Ultra monochrome back-illuminated EMCCD camera, a 2x post-magnification lens and a Borealis Perfect Illumination Delivery™ system. Images acquired with that system have a pixel dimension of 84 × 84 nm and a depth of 16 bits. A single mosaic grid was used to cover each entire brain section with an adaptive *z* focus set-up by the user to follow the unevenness of the tissue using the Andor iQ2 software. In both systems, eGFP was excited using a 488 nm laser and mKO2 with a 561 nm laser. Acquisition parameters were optimized at adult stages when the synapse intensity was high.

**Cohen's *d* formula**

Cohen's *d* values in Fig. 1 measure the effect size of synaptome parameter changes between control and $Pax6^{+/-}$ mice as follows:

$$d = \frac{\overline{x_1} - \overline{x_2}}{s}$$

where $\overline{x_1}$ and $\overline{x_2}$ are the sample average synaptome parameter for the *Pax6*$^{+/-}$ and control groups, respectively, for a given subregion, and $s$ is pooled standard deviation, defined as follows:

$$s = \sqrt{\frac{(n_1 - 1)\, s_1^2 + (n_2 - 1)\, s_2^2}{n_1 + n_2 - 2}}$$

where $s_1$ and $s_2$ are the sample standard deviations in *Pax6*$^{+/-}$ and control groups, respectively, and $n_1$ and $n_2$ are the sample numbers of mutant and control groups, respectively.

The Cohen's *d* values in Fig. 2 and 4 were calculated based on the Bayesian estimation by firstly inferring a probability distribution of Cohen's *d* values. The final Cohen's *d* value was then given as the mode of the distribution. Details of calculation is given in the next section.

**Bayesian analysis**

Bayesian estimation[56] as used previously[14] was also applied to test the significance of the mutant effects on synaptome maps, including subtype density (Fig. 1), similarity matrix (Fig. 2), and synapse diversity (Fig. 4C), and also the difference of SPL mutant effects versus LPL effects (Fig. 4B). The results were finally corrected over all subregions using the Benjamini-Hochberg procedure.

Bayesian estimation was also used in calculating the Cohen's *d* values between *Pax6*$^{+/-}$ and control mice (Figs. 2B,C, 4B,C). Using the Monte Carlo simulation, we upscaled the sample number based on the sample values and a t-distribution model to infer the probability density distributions/functions (PDFs) of the mean and standard deviation of the *Pax6*$^{+/-}$ and control group. Then the distributions (PDFs) of Cohen's *d* values were calculated based on the mean and standard deviation. The final Cohen's *d* was given as the mode of its PDF, namely the Cohen's *d* value that gives the highest probability density.

In Fig. 4 B, we tested whether or not the SPL type has a larger mutant effect size than the LPL in each subregion across all age groups. This was done by comparing the mutant effect size values (Cohen's $d_{mutant}$) of SPL and LPL types using Bayesian analysis. We first inferred the PDFs of mutant effect size, $f_{SPL}(d_{mutant})$ for SPL and $f_{SPL}(d_{mutant})$ for LPL respectively with Monte-carlo simulation. The two PDFs $f_{SPL}(d_{mutant})$ and $f_{SPL}(d_{mutant})$ were then compared using Bayesian analysis to test whether the mutant effect size in SPL is significantly larger or smaller than that in LPL, and by how much using Bayesian estimation of the Cohen's $d_{SPL\text{ - }LPL}$,

$$d_{SPL-LPL} = \frac{E[f_{SPL}(d_{mutant})] - E[f_{LPL}(d_{mutant})]}{\sqrt{(s^2[f_{SPL}(d_{mutant})] + s^2[f_{LPL}(d_{mutant})])/2}}$$

where $E[f]$ and $s[f]$ are, respectively, the expectation and standard deviation of a PDF $f$. The Cohen's $d_{SPL-LPL}$ of SPL against LPL and the corresponding significance *P* values were calculated for each of the 131 subregions in all age groups. The Benjamini-Hochberg multiple comparison correction was finally applied over all subregions to generate the adjusted *P* values.

### Similarity matrices and network analysis

Each row/column in the matrix represents one delineated brain subregion at one age of either control or mutant group (Figs. 2A and S3). Elements in the matrix are the synaptome similarities between two subregions quantified by differences in standardized synaptome parameters. The similarity ratio (Sratio) of the matrix in the *Pax6*$^{+/-}$ and control mice of different ages in Fig. S3 is calculated as the similarity of two subregions from different main regions (corresponding to areas outside white boxes distributed diagonally in the similarity matrices in Figs. 2A and S4) divided by similarity of two subregions in the same main region (the areas marked by the white boxes lying on the diagonal in Figs. 2A and S4). A high $S_{ratio}$ value indicates a homogeneous synaptome similarity distributed over all columns/rows in the matrix, whereas a low ratio indicates homogeneous synaptome similarity only within the same main region. Details concerning the calculation of similarity matrix and ratio can be found in previous work[13,14].

After the ratio $S_{ratio}$ is calculated for each subregion in the *Pax6*$^{+/-}$ and control mice of different ages, the Bayesian estimation was used to give a developmental trajectory of Cohens' *d* with P values in Fig. 2B.

The network analysis was based on the similarity matrices of individual brain sections quantified in a similar way to those in Figs. 2A and S4. Nodes in the network are representations of the delineated subregion. The small worldness is the topology quantification of the network, where the whole set of nodes is divided into small and clustered groups: nodes within the same groups are highly connected/similar, whereas those between groups are disconnected/dissimilar[57]. Details of the small-worldness calculation can be found in our previous studies[13,14]. With the small-worldness values being calculated for each of all ages in the *Pax*6$^{+/-}$ and control mice, the Bayesian test was used to calculate the Cohen's *d* and *P* values in Fig. 2C.

**Computational modeling of synaptic responses**

Computational modeling of synaptic responses was based on our previously described models[13,14] representing physiology at several age points.

Synaptic gradients along the radial and tangential directions (Fig 3) are represented with gradient intensity symbolized by color intensity; PSD95 (green) and SAP102 (magenta) were used as in previous work[13,14] to set synaptic properties of the computational model. To model synaptic physiology corresponding to P1, P7, P28, P35 and P56 in control and *Pax6*$^{+/-}$ -animals, differences in intensity of PSD95 and SAP102 along radial as well as tangential directions of CA1 sr were computed.

Computing gradients. For each of the age groups, we first computed the geometric mean (control N=5, *Pax6*$^{+/-}$ N=4) of intensity of PSD95 and SAP102 for delineations of CA1sr into four radial and ten tangential subregions. Next, we computed the mean of each radial and tangential direction and then normalized this by the min and max values among all time points for control and *Pax6*$^{+/-}$. Thus, each control and *Pax6*$^{+/-}$, PSD95 and SAP102, radial/tangential value expresses the relative intensity compared over all time points and regardless of genotype. This relative intensity was used to scale the spatial gradients used in previous work[14]. This scaling thus takes into consideration the observation[13] that intensities of PSD95 and SAP102 increase from P1 and attain a maximum at 3 months, which is the age of the study by Zhu et al.[14], and allows for a comparison of *Pax6*$^{+/-}$ versus control relative intensity levels over the age points studied here.

In the model, weighted gradients described above were used to scale the amplitude of short-term depression and facilitation (the PSD95 and SAP102 tangential gradient decrement factor, respectively), as well as the time constant of short-term depression and facilitation (the PSD95 and SAP102 radial gradient decrement factor, respectively). Our model is constrained to consider only age-dependent and genotype-dependent changes in synapse protein composition and does not consider potential changes in dendritic morphology or other neuronal properties. Synaptic responses following a stimulation pattern were first quantified for each synapse as the sum of synaptic max amplitudes reached following each of the 20 stimulus pulses. Differences in synaptic responses between control and *Pax6*$^{+/-}$ mice of all age groups and all stimulation patterns were assessed using a paired t-test with corrections for multiple comparisons using Benjamini-Hochberg correction, where all the 121 synaptic responses of the control case were compared with those from the *Pax6*$^{+/-}$ case.

### Supplementary Figures

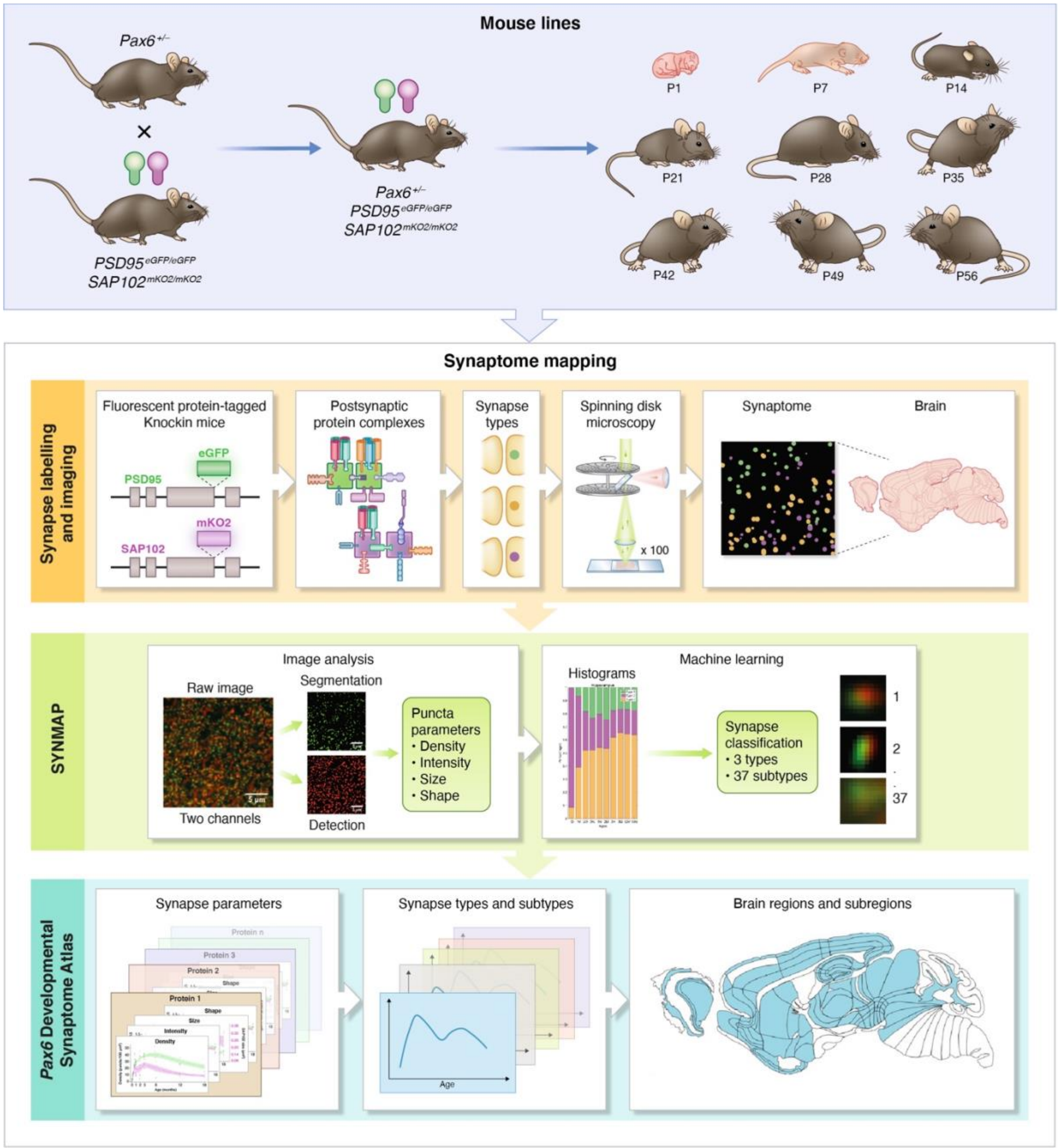


**Figure S1. Procedure for mapping the synaptome architecture of *Pax6*$^{+/-}$ mice.**
Mouse lines: Mice carrying the heterozygous *Pax6* mutation were crossed with mice carrying the fluorescent-tagged PSD95 and SAP102 proteins to produce cohorts of

*Pax6*$^{+/-}$;*Psd95*$^{eGFP/eGFP}$;*Sap102*$^{mKO2/mKO2}$ and control *Pax6*$^{+/+}$;*Psd95*$^{eGFP/eGFP}$;*Sap102*$^{mKO2/mKO2}$ mice.

Synaptic labeling and imaging: The first panel illustrates the genetic tagging of the endogenous *Psd95* and *Sap102* loci with eGFP and mKO2 fluorescent proteins. These genetic modifications result in expression of PSD95-eGFP and SAP102-mKO2 fusion proteins that assemble with ion channels and other proteins into postsynaptic protein complexes. The differential distribution of these proteins into synapses produces synapse type diversity, which can be visualized with a spinning disk confocal microscope in brain tissue sections.

SYNMAP: Raw images of fluorescent synaptic puncta are detected, segmented and the density, intensity, size and shape of each punctum determined. The puncta are then classified into 3 types and 37 subtypes based on these molecular and morphological parameters.

Pax6 Developmental Dynaptome Atlas: The synaptic parameters, density of synapse types and subtypes are quantified in 131 subregions and their spatial maps and temporal trajectories determined. The Pax6 Developmental Synaptome Atlas is available online[41].

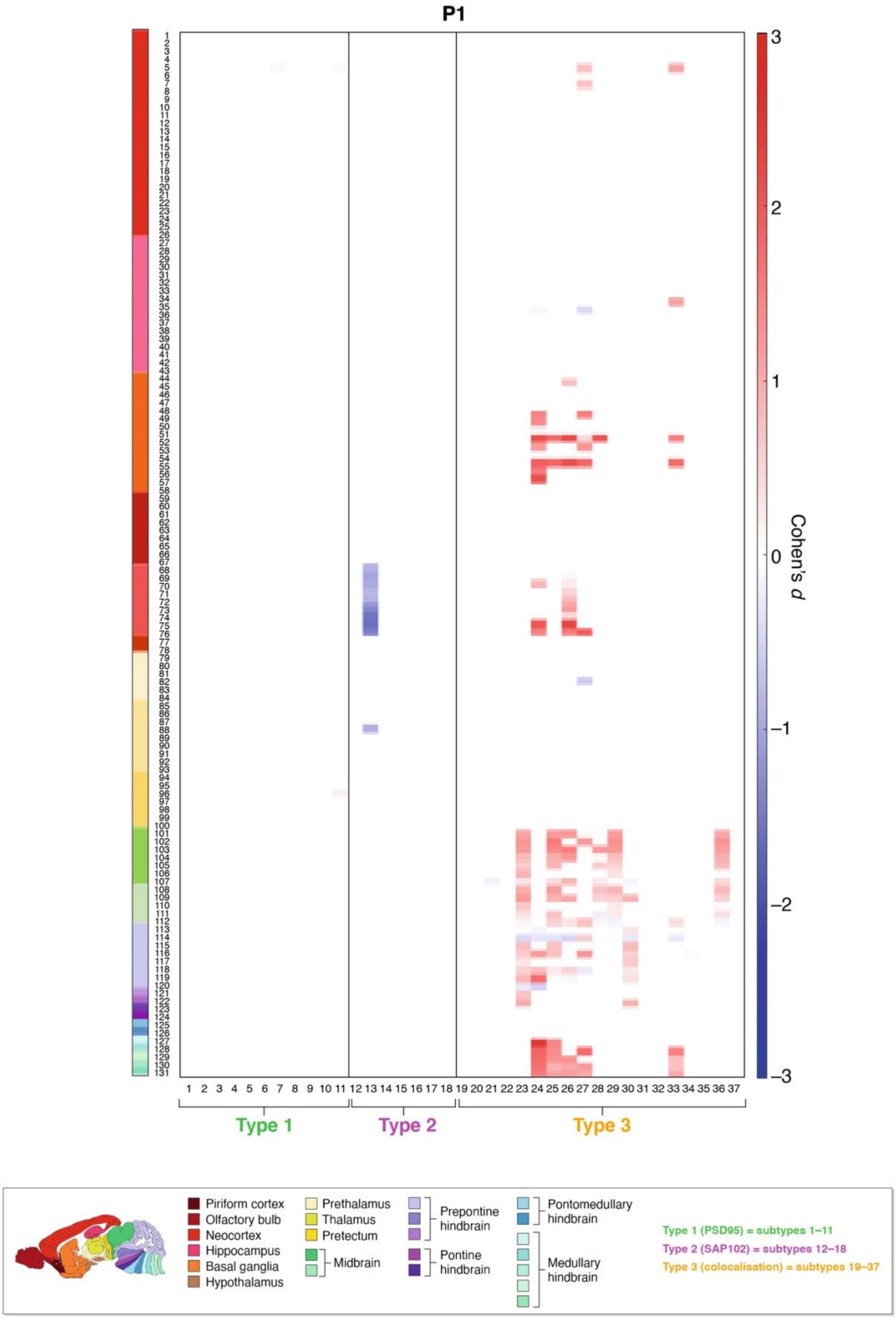
P1
Cohen's d
Type 1
Type 2
Type 3
Piriform cortex
Olfactory bulb
Neocortex
Hippocampus
Basal ganglia
Hypothalamus
Prethalamus
Thalamus
Pretectum
Midbrain
Prepontine hindbrain
Pontine hindbrain
Pontomedullary hindbrain
Medullary hindbrain
Type 1 (PSD95) = subtypes 1–11
Type 2 (SAP102) = subtypes 12–18
Type 3 (colocalisation) = subtypes 19–37

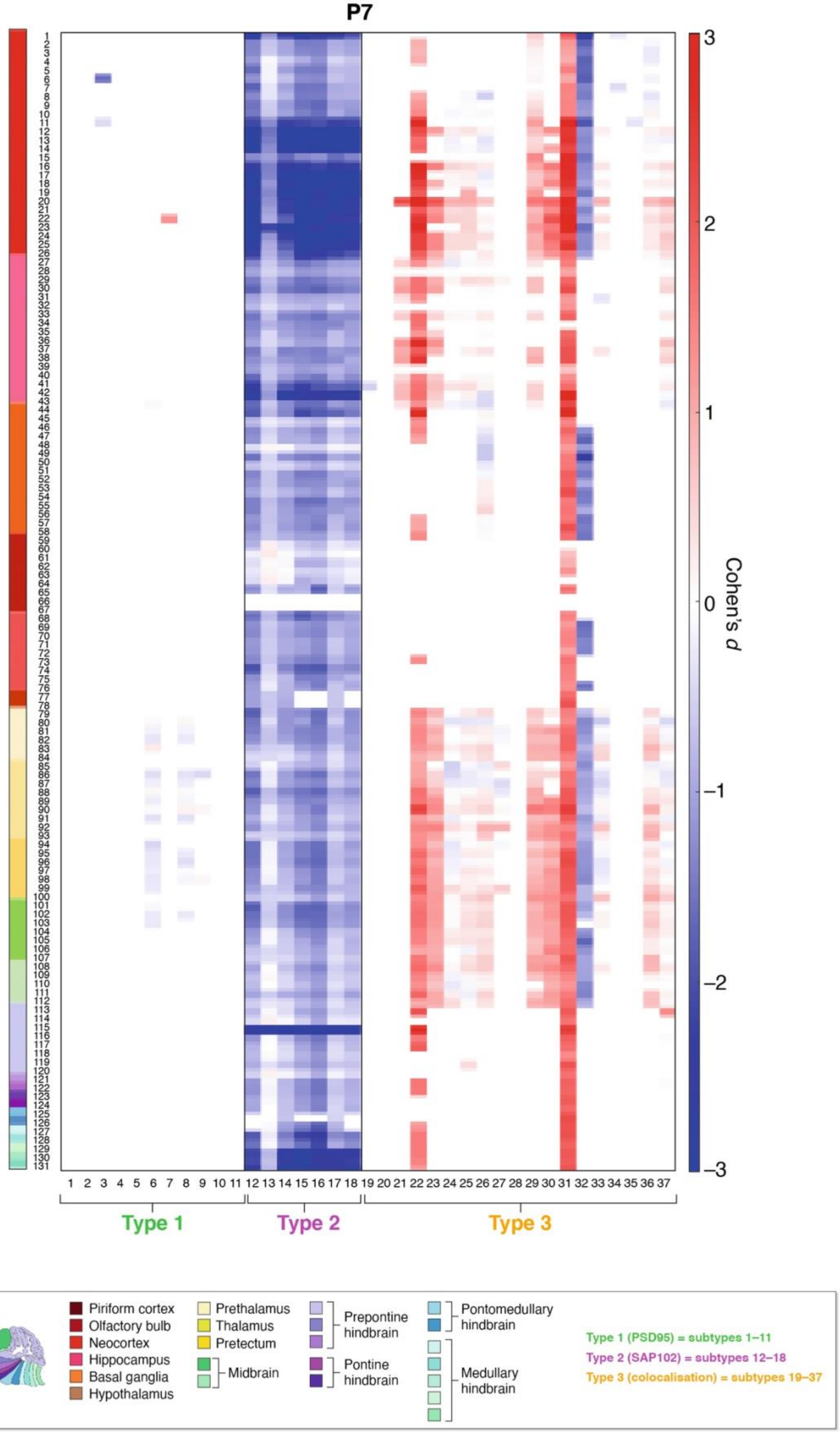
P7
Cohen's d
Type 1
Type 2
Type 3
Piriform cortex
Olfactory bulb
Neocortex
Hippocampus
Basal ganglia
Hypothalamus
Prethalamus
Thalamus
Pretectum
Midbrain
Prepontine hindbrain
Pontine hindbrain
Pontomedullary hindbrain
Medullary hindbrain
Type 1 (PSD95) = subtypes 1–11
Type 2 (SAP102) = subtypes 12–18
Type 3 (colocalisation) = subtypes 19–37

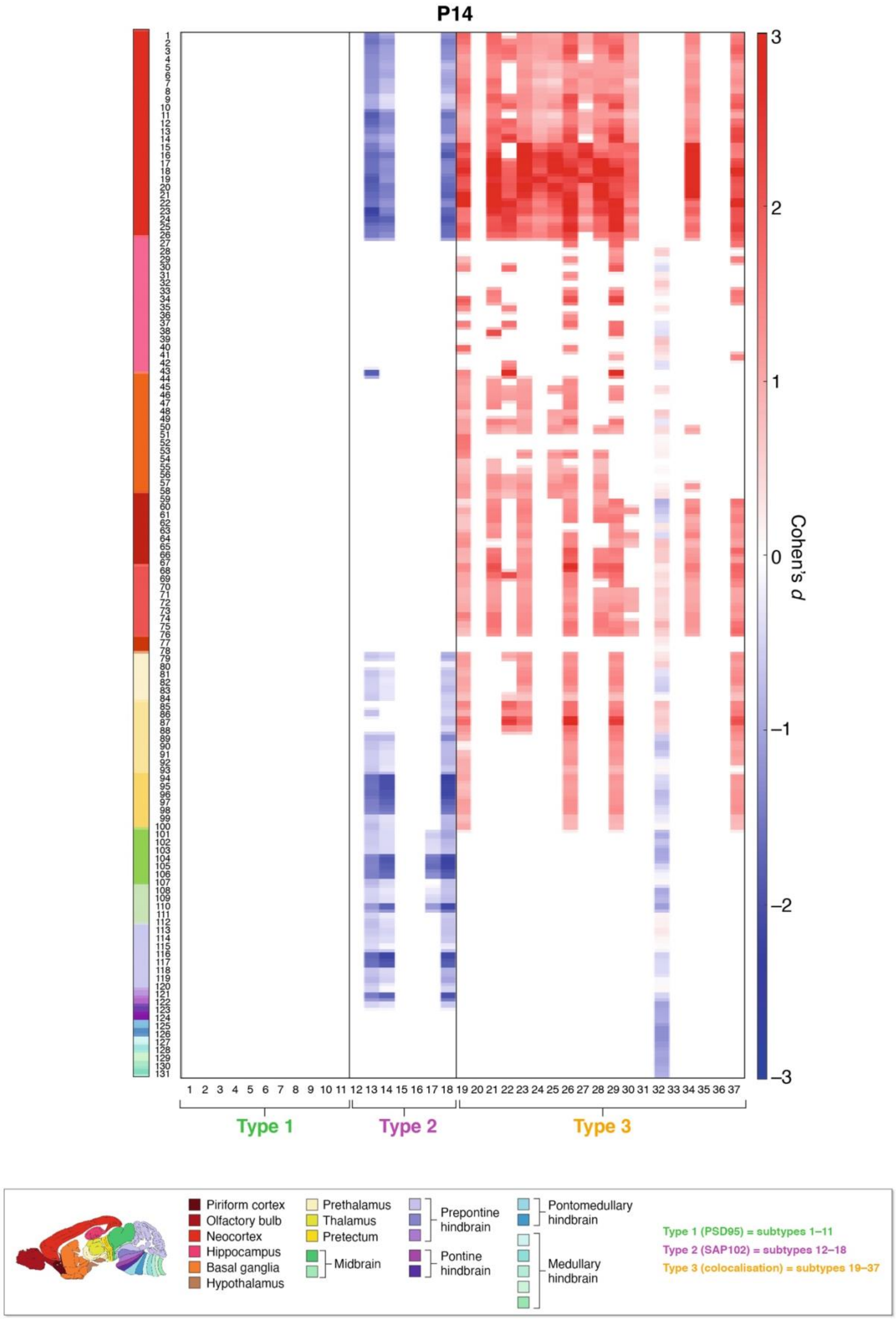
P14
Cohen's d
3
2
1
0
−1
−2
−3
1 2 3 4 5 6 7 8 9 10 11 12 13 14 15 16 17 18 19 20 21 22 23 24 25 26 27 28 29 30 31 32 33 34 35 36 37
Type 1
Type 2
Type 3
Piriform cortex
Olfactory bulb
Neocortex
Hippocampus
Basal ganglia
Hypothalamus
Prethalamus
Thalamus
Pretectum
Midbrain
Prepontine hindbrain
Pontine hindbrain
Pontomedullary hindbrain
Medullary hindbrain
Type 1 (PSD95) = subtypes 1–11
Type 2 (SAP102) = subtypes 12–18
Type 3 (colocalisation) = subtypes 19–37

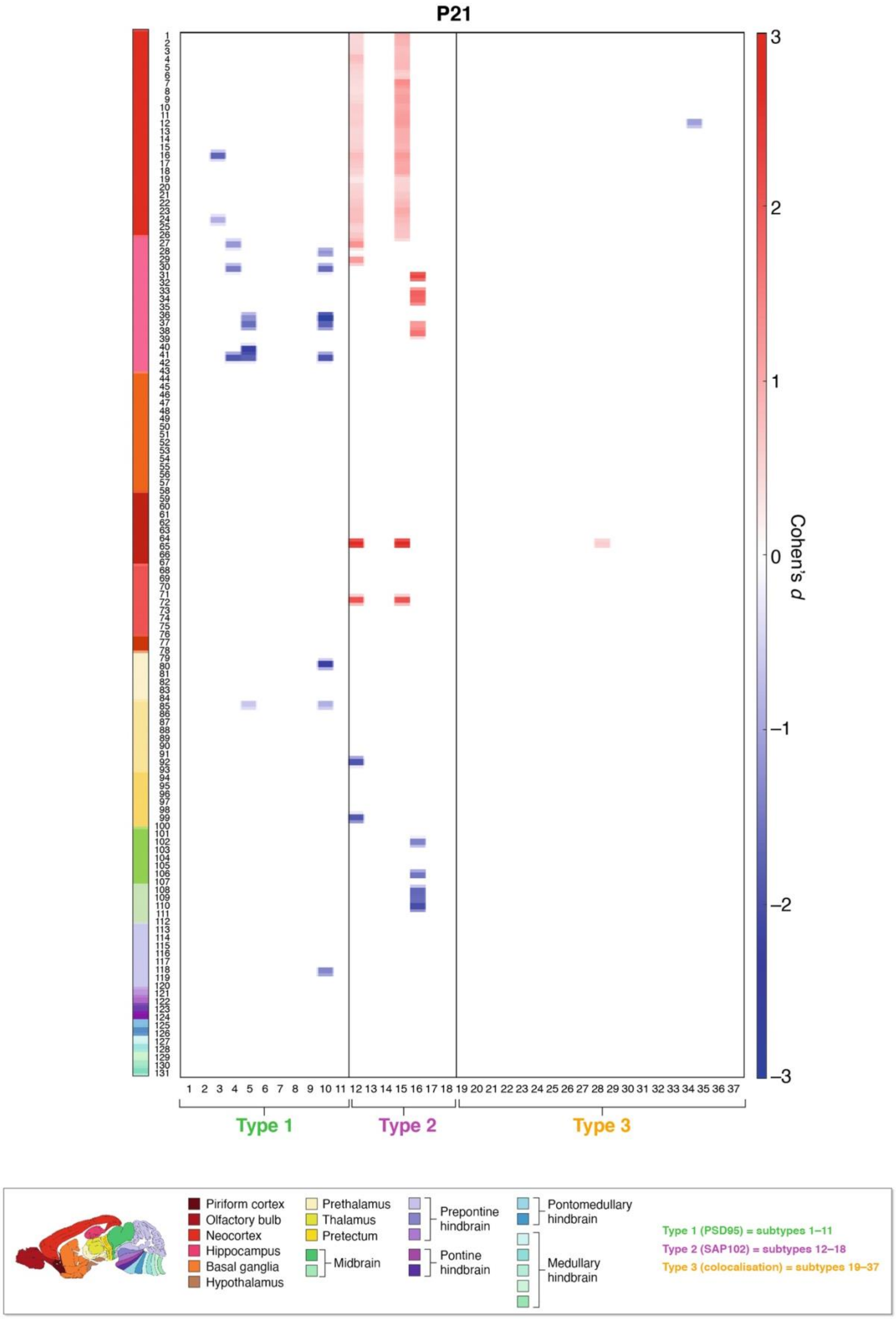
P21
Cohen's d
3
2
1
0
−1
−2
−3
Type 1
Type 2
Type 3
Piriform cortex
Olfactory bulb
Neocortex
Hippocampus
Basal ganglia
Hypothalamus
Prethalamus
Thalamus
Pretectum
Midbrain
Prepontine hindbrain
Pontine hindbrain
Pontomedullary hindbrain
Medullary hindbrain
Type 1 (PSD95) = subtypes 1–11
Type 2 (SAP102) = subtypes 12–18
Type 3 (colocalisation) = subtypes 19–37

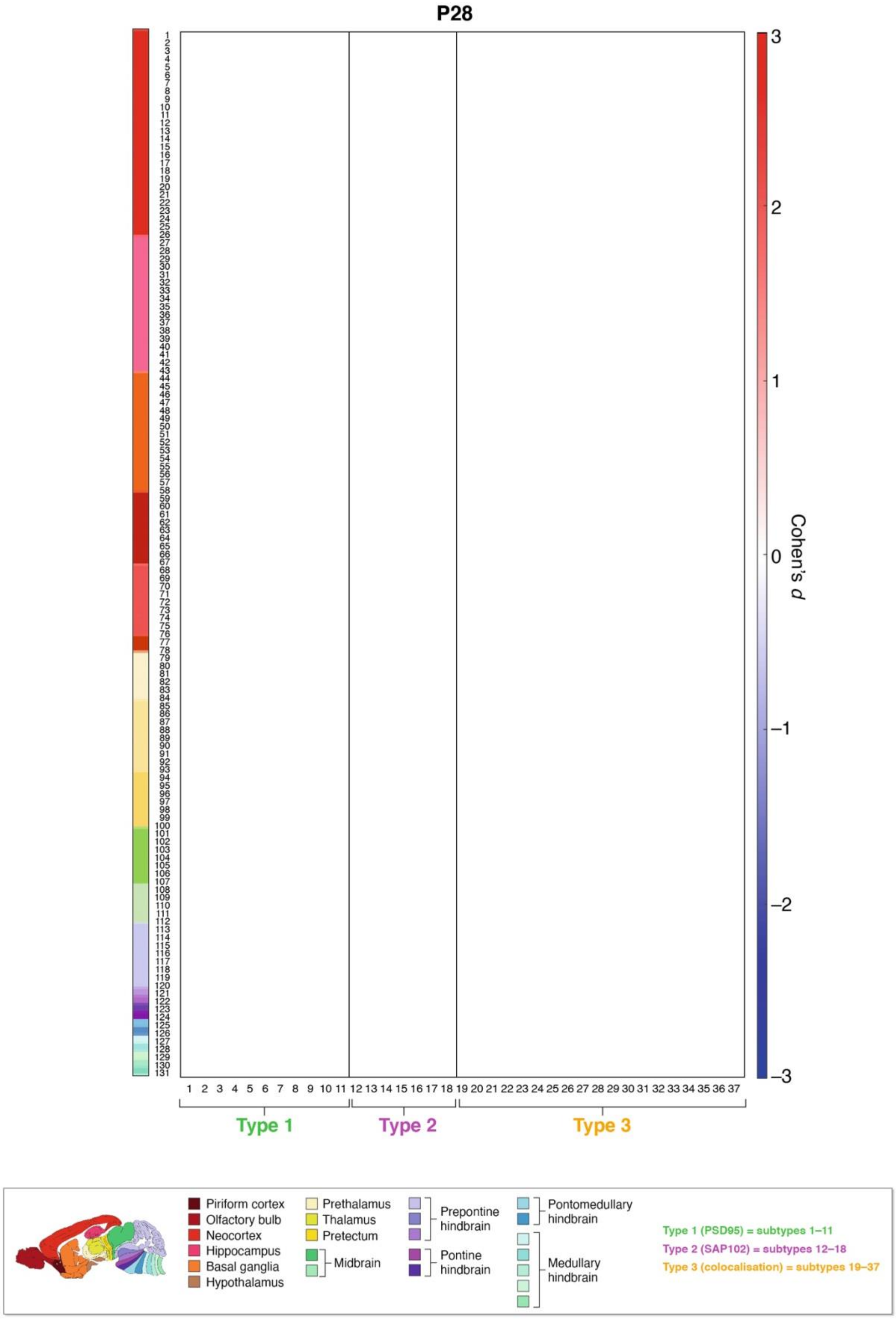

P28
Cohen's d
Type 1
Type 2
Type 3
Piriform cortex
Olfactory bulb
Neocortex
Hippocampus
Basal ganglia
Hypothalamus
Prethalamus
Thalamus
Pretectum
Midbrain
Prepontine hindbrain
Pontine hindbrain
Pontomedullary hindbrain
Medullary hindbrain
Type 1 (PSD95) = subtypes 1–11
Type 2 (SAP102) = subtypes 12–18
Type 3 (colocalisation) = subtypes 19–37

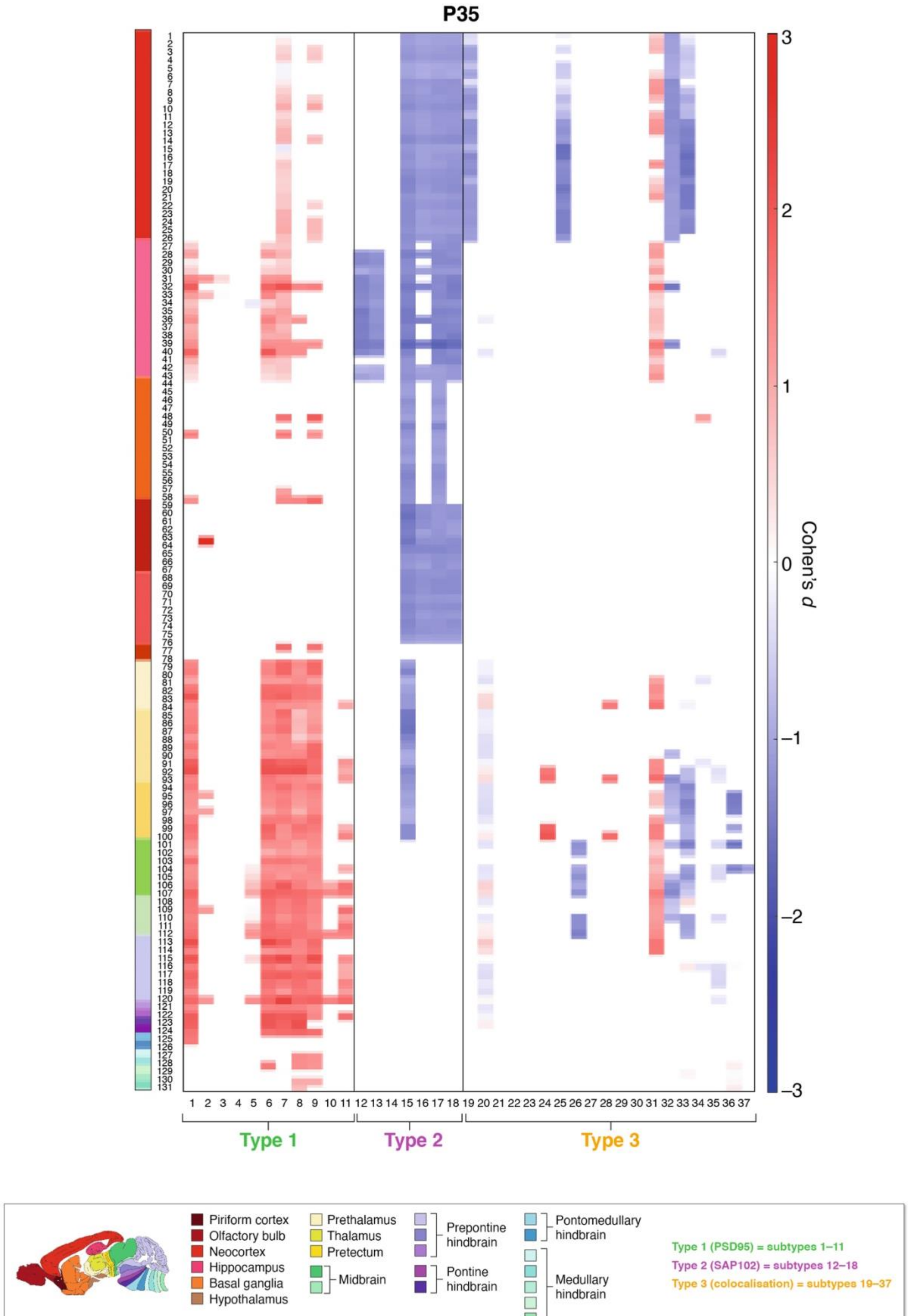

P35
Cohen's d
Type 1
Type 2
Type 3
Piriform cortex
Olfactory bulb
Neocortex
Hippocampus
Basal ganglia
Hypothalamus
Prethalamus
Thalamus
Pretectum
Midbrain
Prepontine hindbrain
Pontine hindbrain
Pontomedullary hindbrain
Medullary hindbrain
Type 1 (PSD95) = subtypes 1–11
Type 2 (SAP102) = subtypes 12–18
Type 3 (colocalisation) = subtypes 19–37

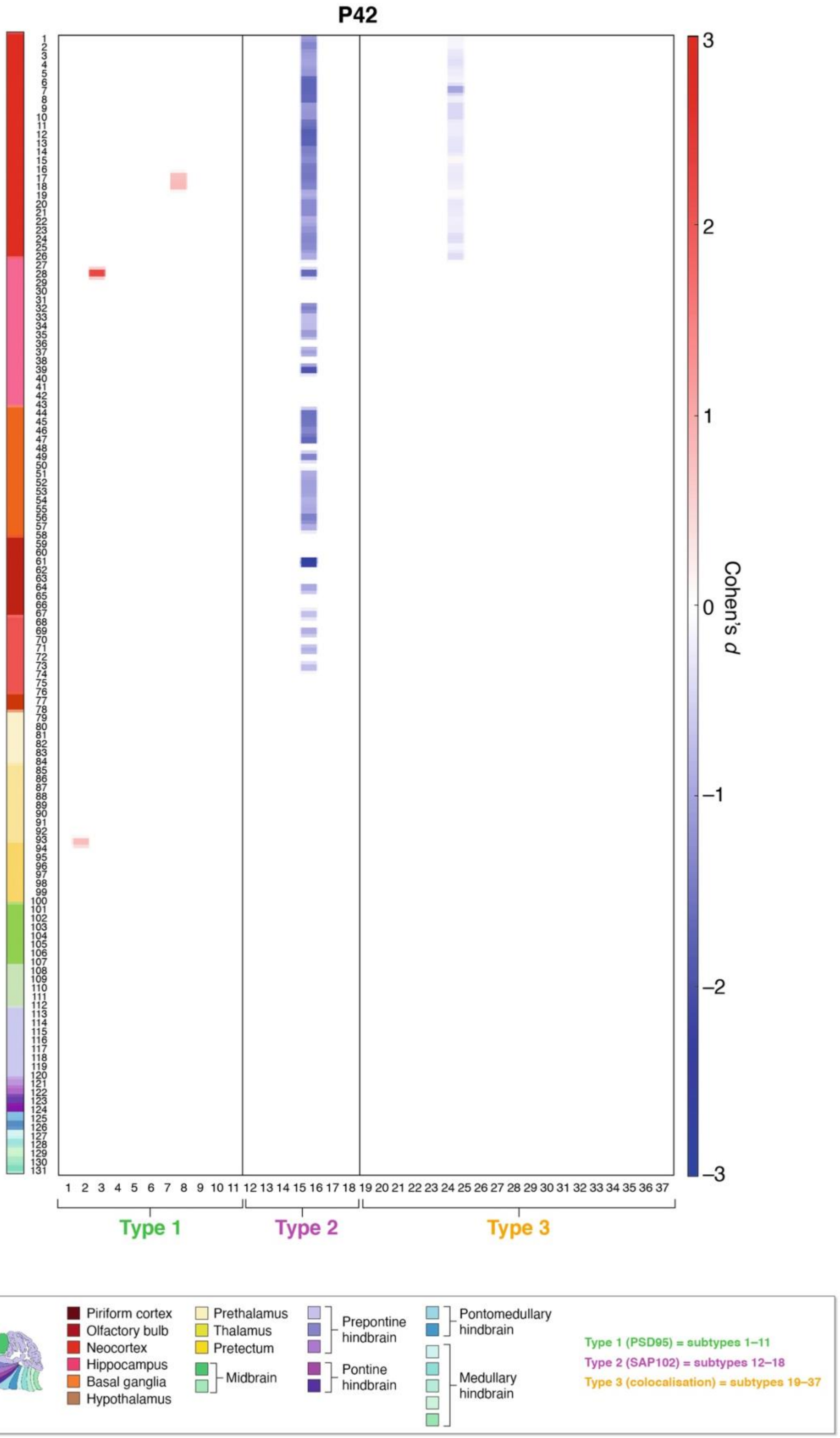
P42
Cohen's d
3
2
1
0
−1
−2
−3
Type 1
Type 2
Type 3
Piriform cortex
Olfactory bulb
Neocortex
Hippocampus
Basal ganglia
Hypothalamus
Prethalamus
Thalamus
Pretectum
Midbrain
Prepontine hindbrain
Pontine hindbrain
Pontomedullary hindbrain
Medullary hindbrain
Type 1 (PSD95) = subtypes 1–11
Type 2 (SAP102) = subtypes 12–18
Type 3 (colocalisation) = subtypes 19–37

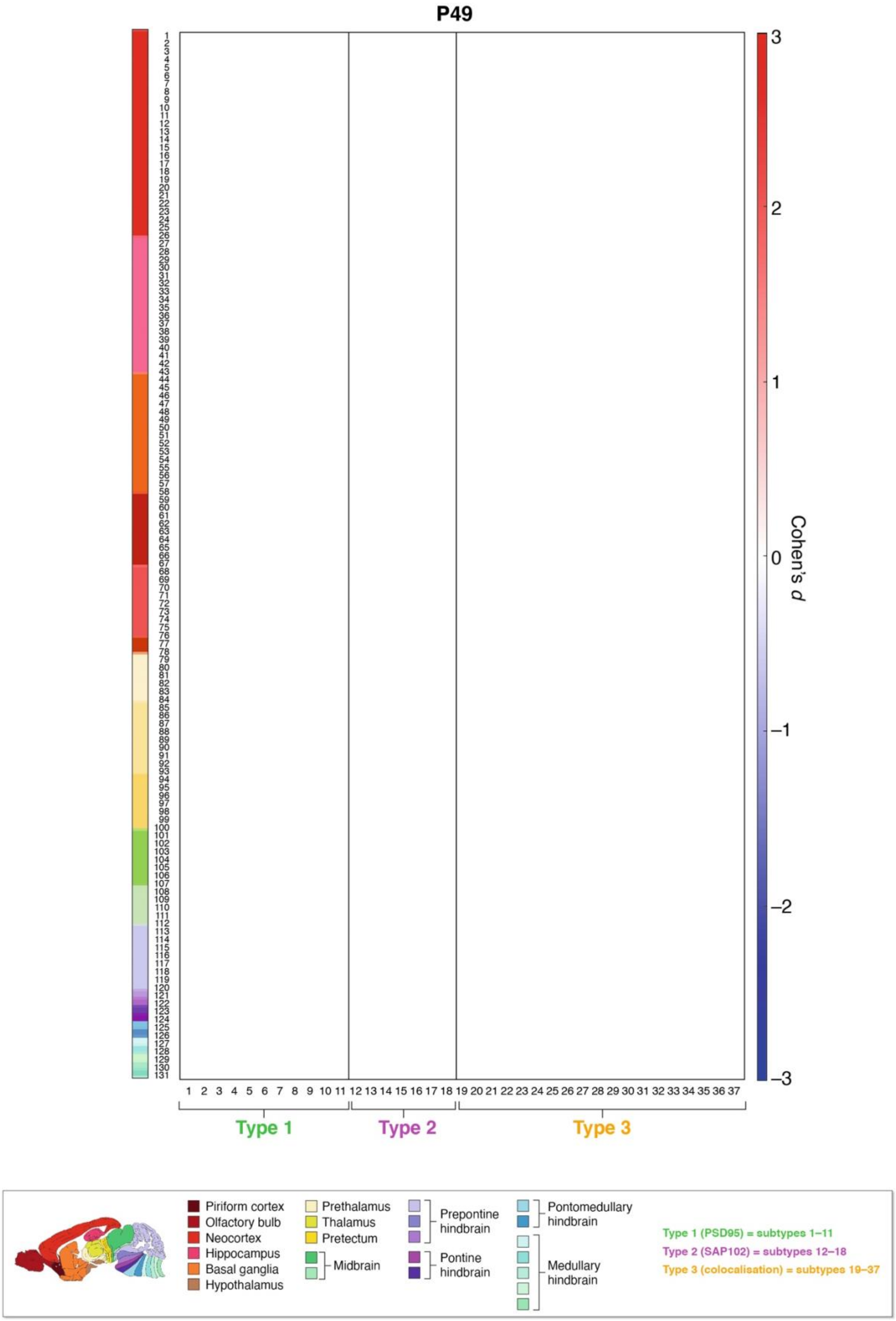
P49
Cohen's d
3
2
1
0
−1
−2
−3
Type 1
Type 2
Type 3
Piriform cortex
Olfactory bulb
Neocortex
Hippocampus
Basal ganglia
Hypothalamus
Prethalamus
Thalamus
Pretectum
Midbrain
Prepontine hindbrain
Pontine hindbrain
Pontomedullary hindbrain
Medullary hindbrain
Type 1 (PSD95) = subtypes 1–11
Type 2 (SAP102) = subtypes 12–18
Type 3 (colocalisation) = subtypes 19–37

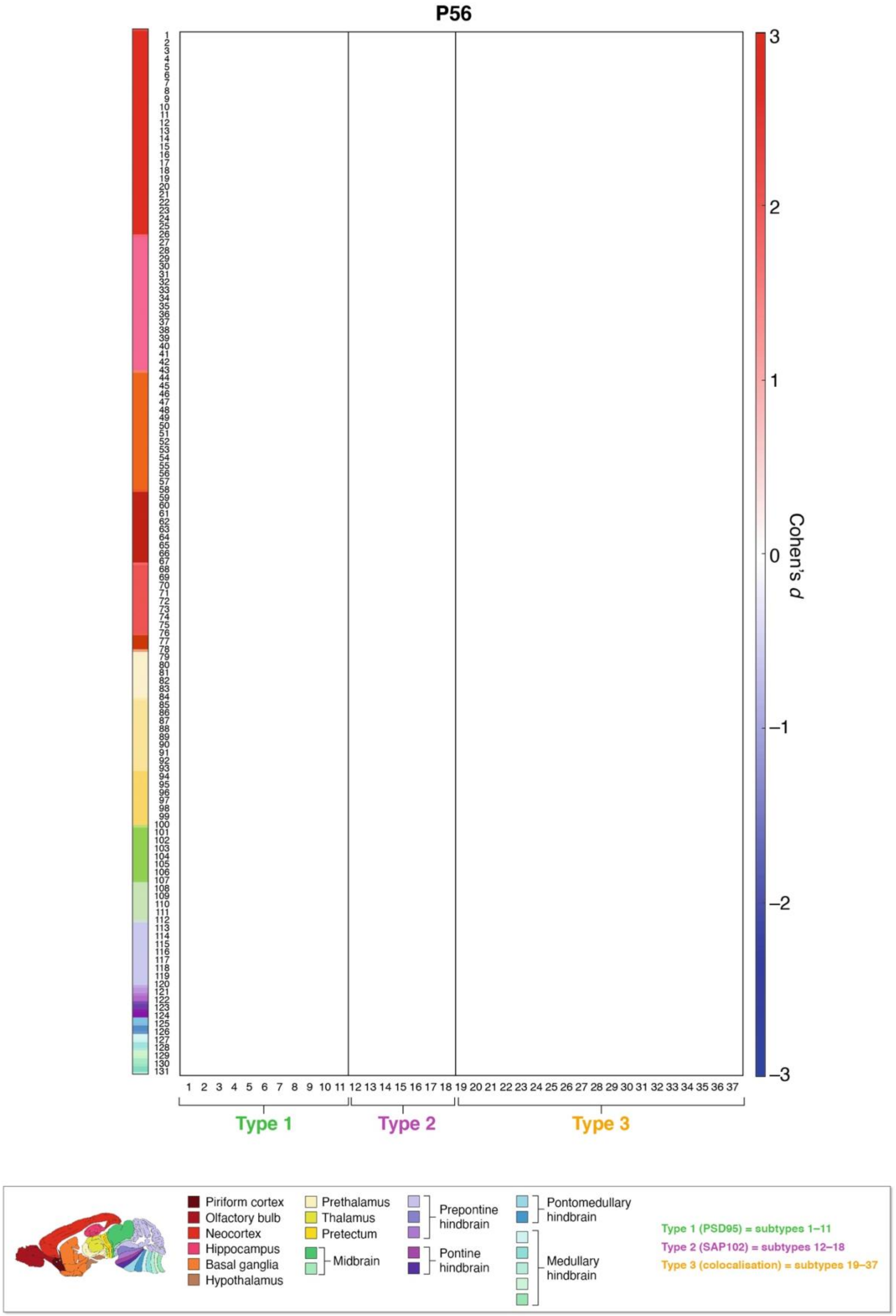
P56
Cohen's d
3
2
1
0
−1
−2
−3
Type 1
Type 2
Type 3
Piriform cortex
Olfactory bulb
Neocortex
Hippocampus
Basal ganglia
Hypothalamus
Prethalamus
Thalamus
Pretectum
Midbrain
Prepontine hindbrain
Pontine hindbrain
Pontomedullary hindbrain
Medullary hindbrain
Type 1 (PSD95) = subtypes 1–11
Type 2 (SAP102) = subtypes 12–18
Type 3 (colocalisation) = subtypes 19–37

**Figure S2. Synaptome phenotypes in *Pax6*$^{+/-}$ mice.**

The difference (Cohen's *d*) in synapse type and subtype density in 131 brain subregions at nine ages between birth and maturity (P1, P7, P14, P21, P28, P35, P42, P49, P56) between *Pax6*$^{+/-}$ and control mice. Significantly different ($P < 0.05$, Benjamini-Hochberg corrected) subregions are shown.

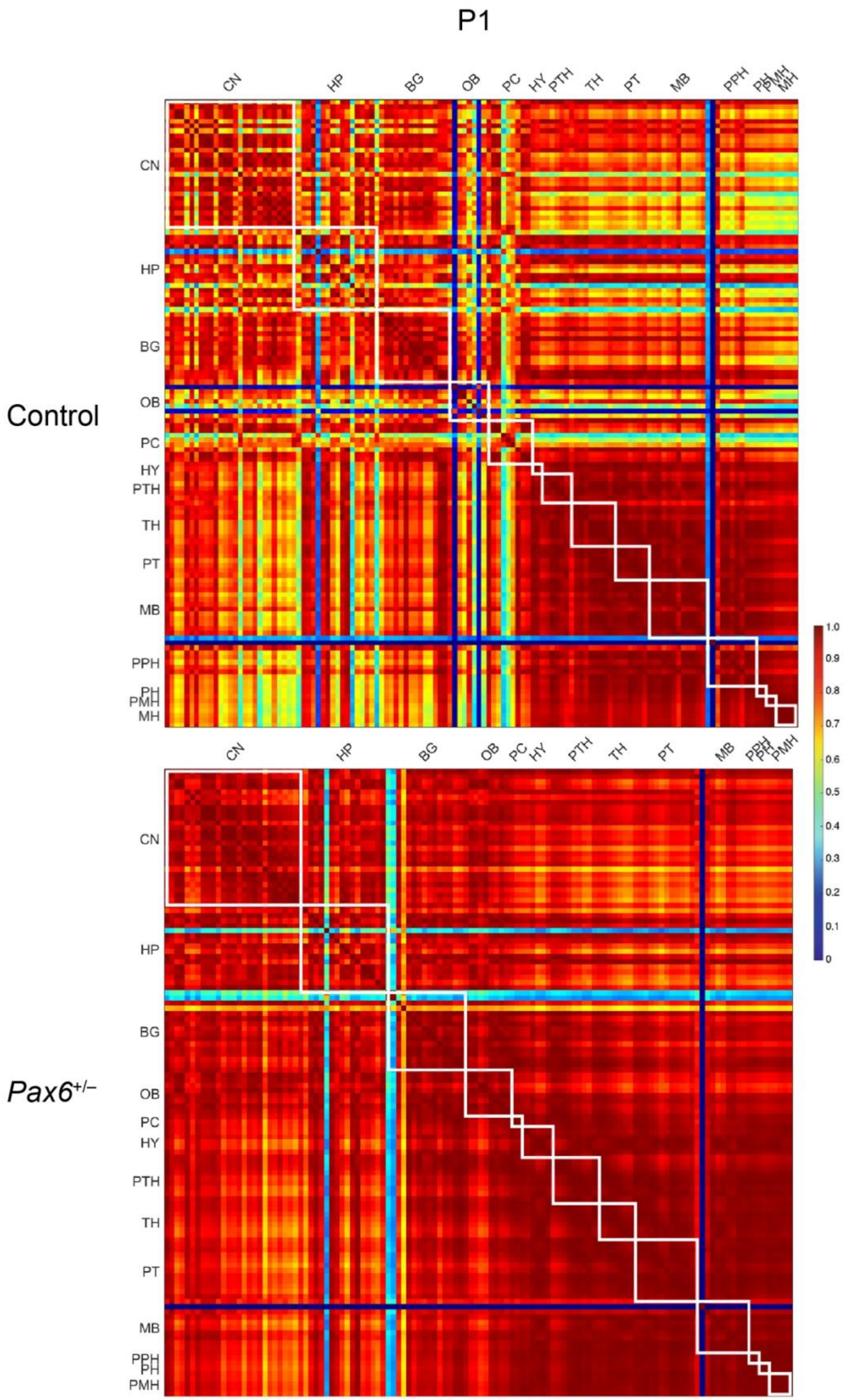
P1
Control
Pax6+/−
CN
HP
BG
OB
PC
HY
PTH
TH
PT
MB
PPH
PH
PMH
MH
1.0
0.9
0.8
0.7
0.6
0.5
0.4
0.3
0.2
0.1
0

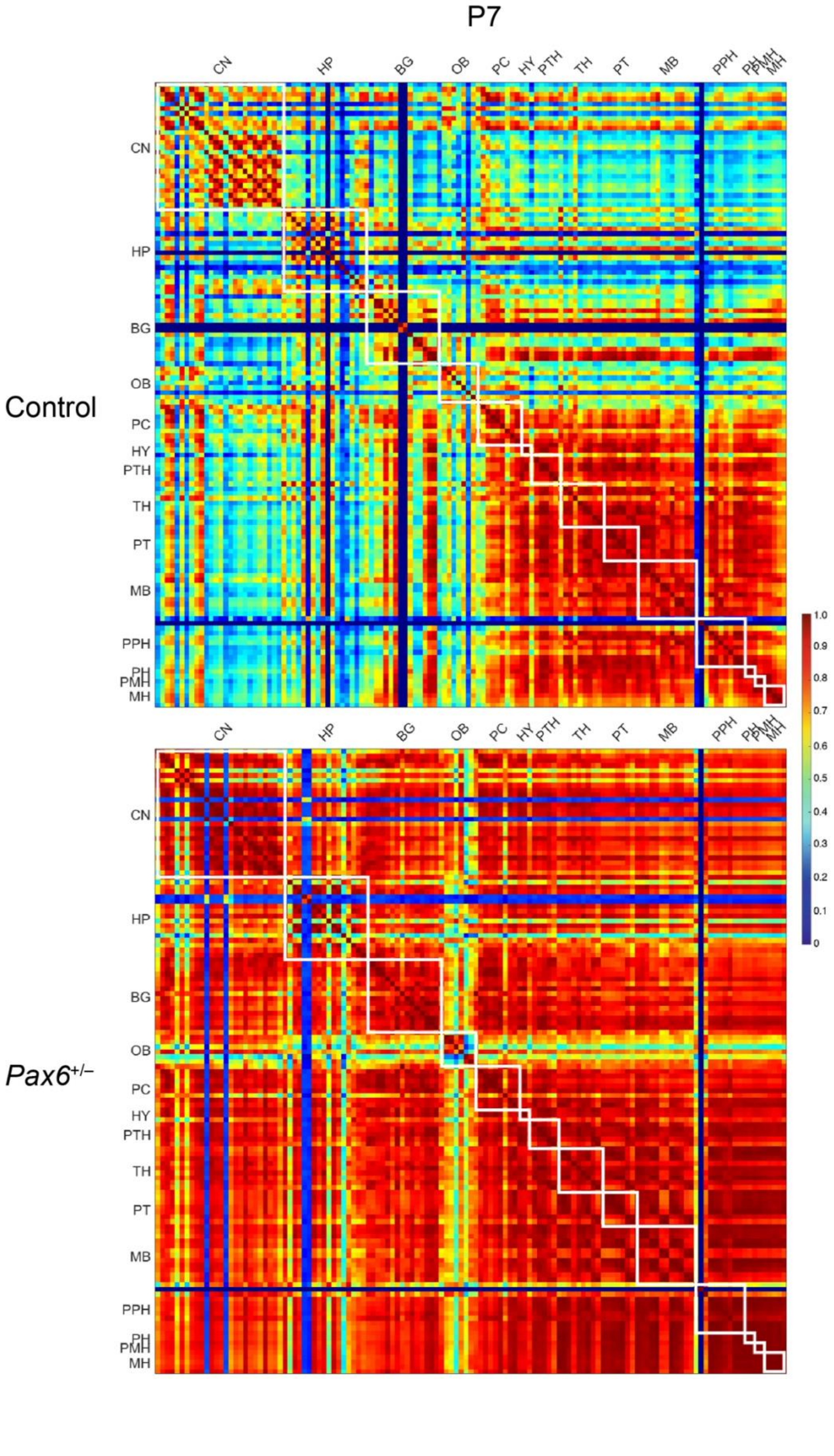
P7
Control
Pax6+/–
CN
HP
BG
OB
PC
HY
PTH
TH
PT
MB
PPH
PH
PMH
MH
1.0
0.9
0.8
0.7
0.6
0.5
0.4
0.3
0.2
0.1
0

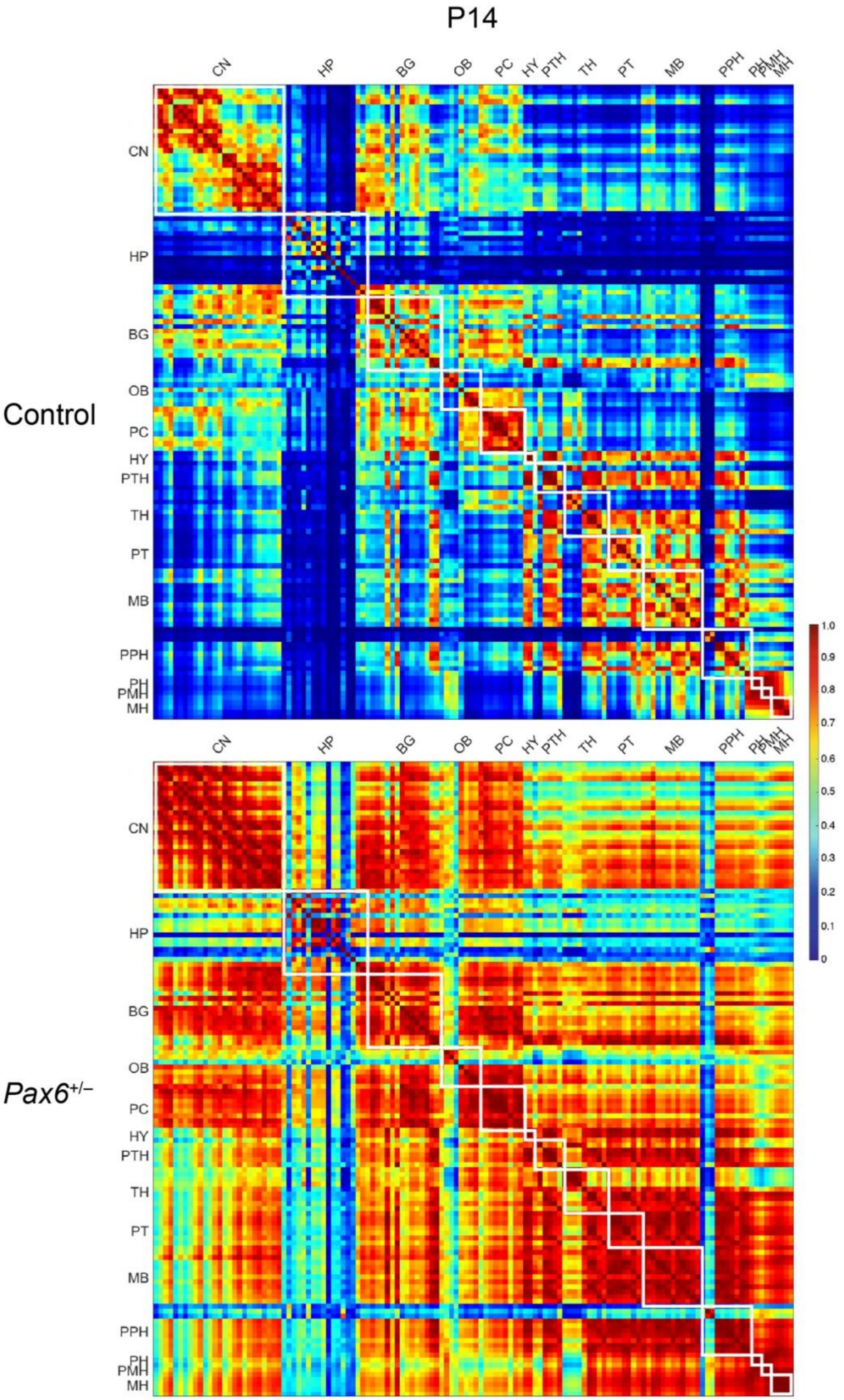

P14
Control
Pax6+/−
CN
HP
BG
OB
PC
HY
PTH
TH
PT
MB
PPH
PH
PMH
MH
1.0
0.9
0.8
0.7
0.6
0.5
0.4
0.3
0.2
0.1
0

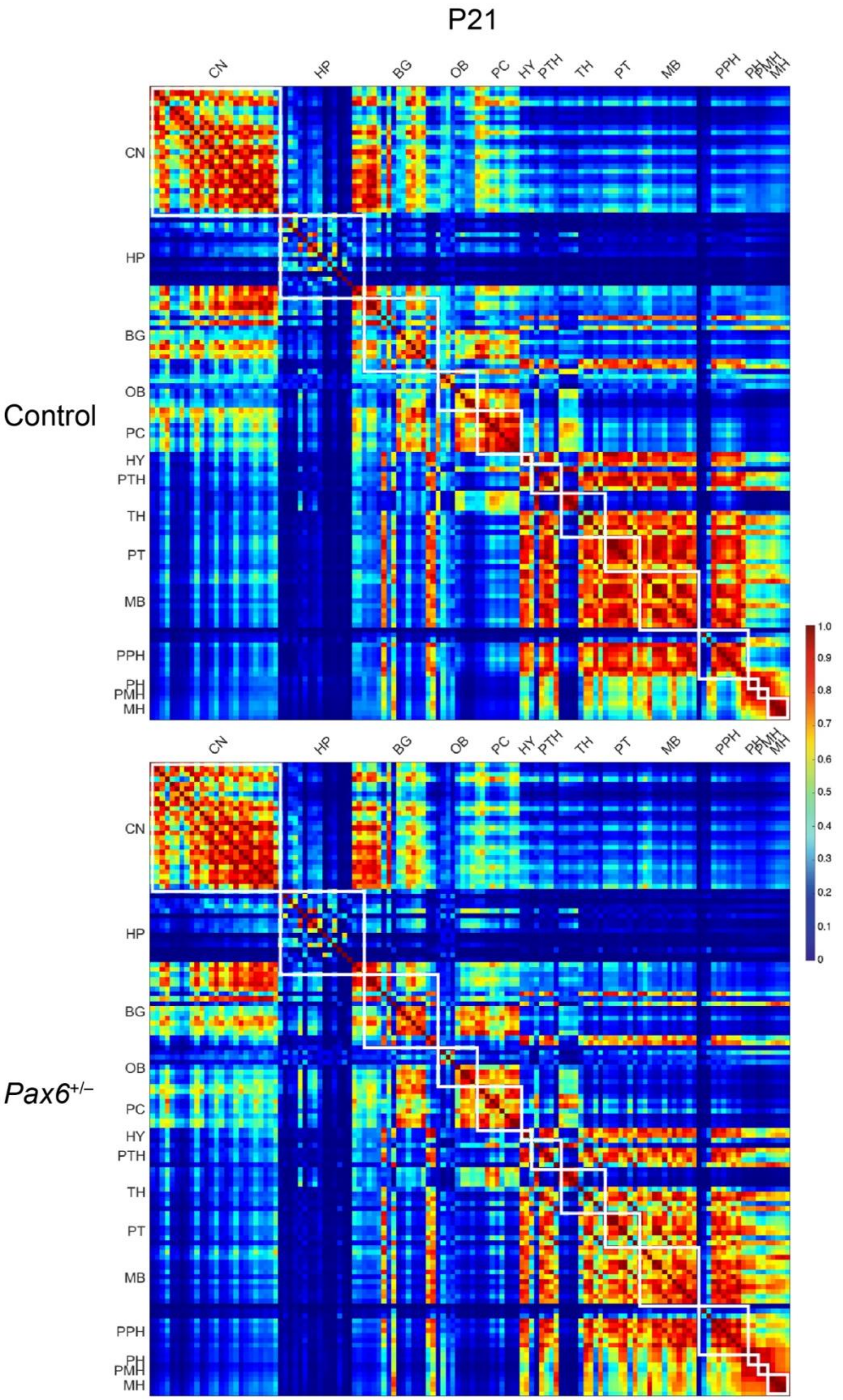
P21
Control
Pax6+/−
CN
HP
BG
OB
PC
HY
PTH
TH
PT
MB
PPH
PH
PMH
MH
1.0
0.9
0.8
0.7
0.6
0.5
0.4
0.3
0.2
0.1
0

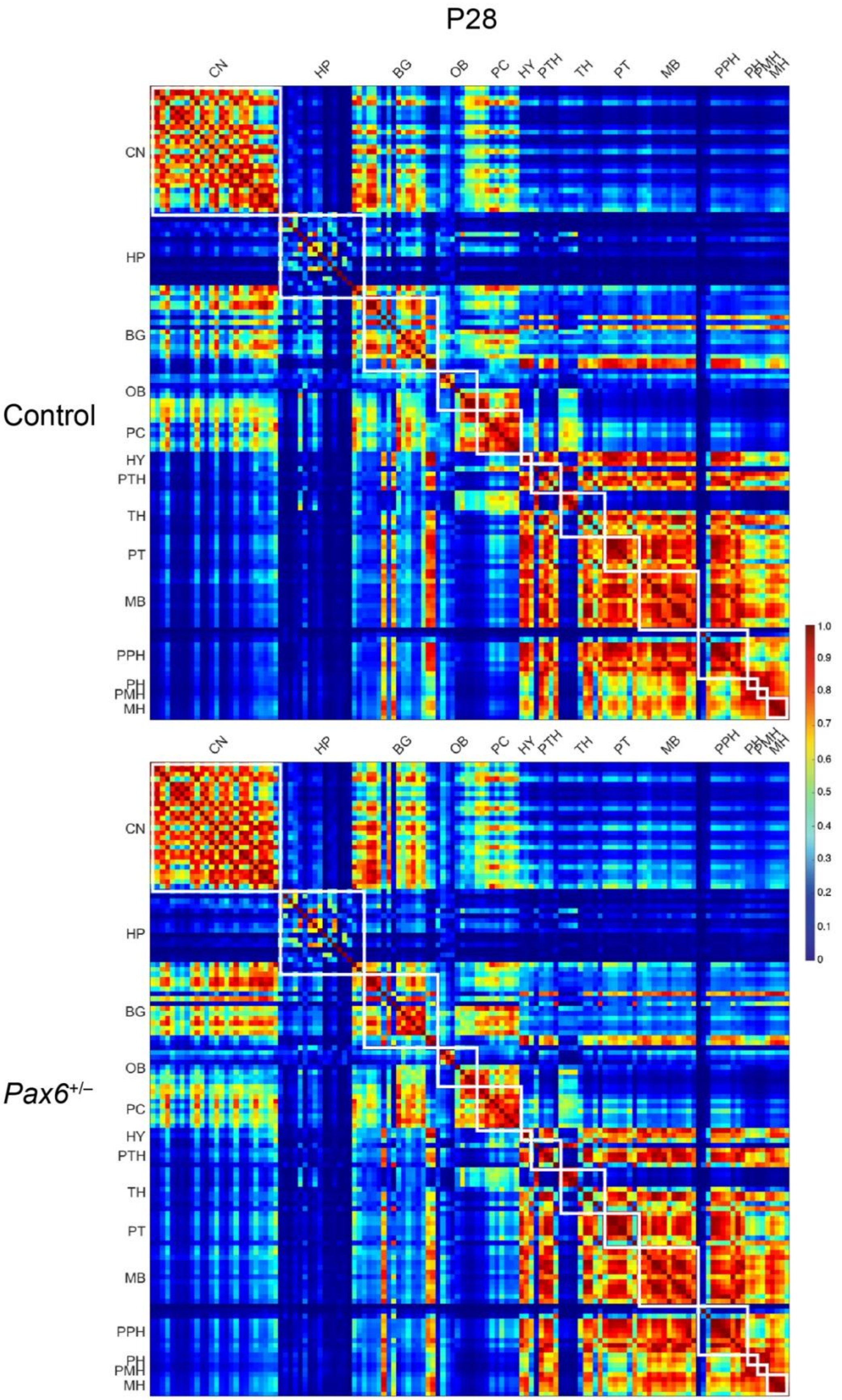
P28
Control
Pax6+/−
CN
HP
BG
OB
PC
HY
PTH
TH
PT
MB
PPH
PH
PMH
MH
1.0
0.9
0.8
0.7
0.6
0.5
0.4
0.3
0.2
0.1
0

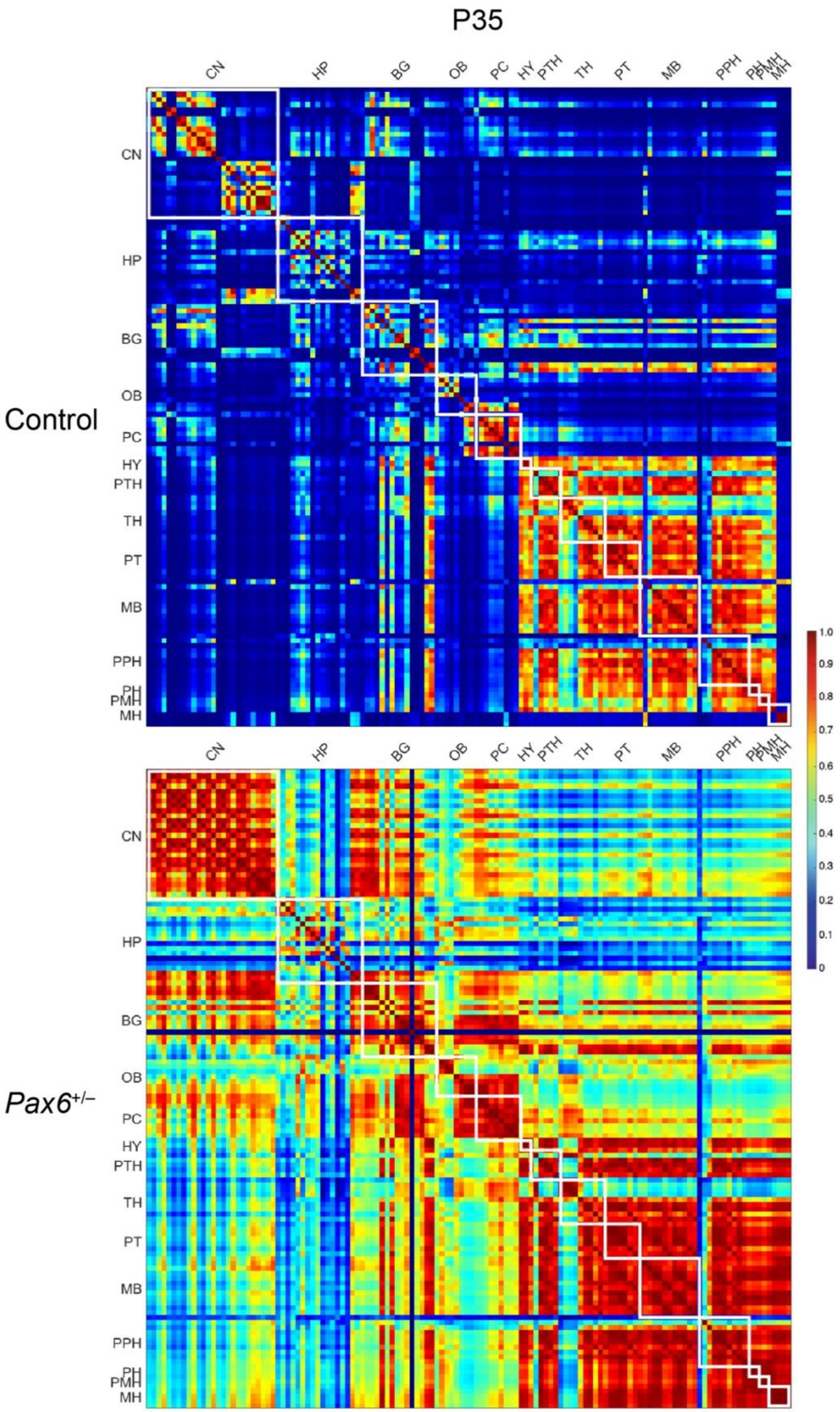
P35
Control
Pax6+/–
CN
HP
BG
OB
PC
HY
PTH
TH
PT
MB
PPH
PH
PMH
MH
1.0
0.9
0.8
0.7
0.6
0.5
0.4
0.3
0.2
0.1
0

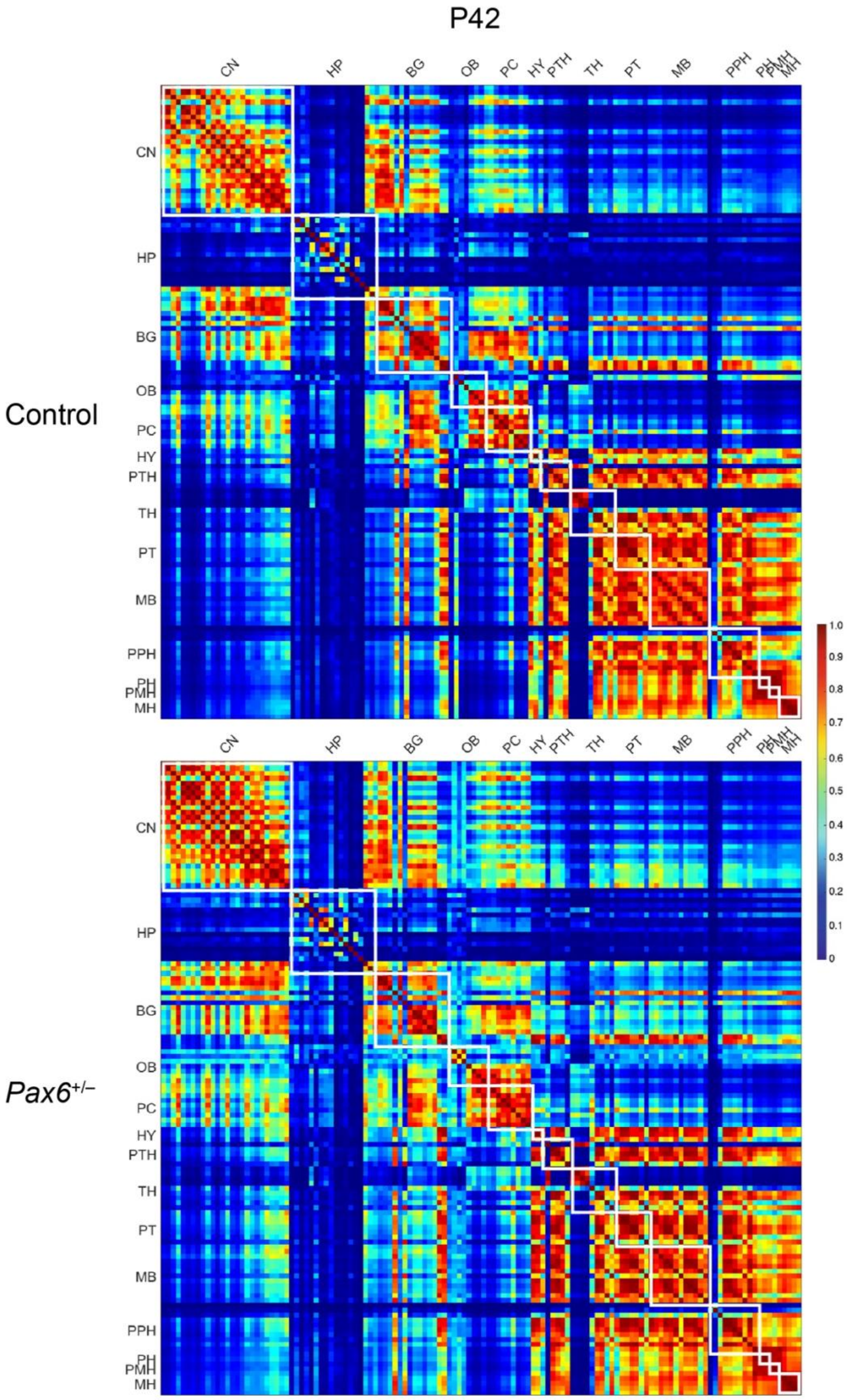
P42
Control
Pax6+/−
CN
HP
BG
OB
PC
HY
PTH
TH
PT
MB
PPH
PH
PMH
MH
1.0
0.9
0.8
0.7
0.6
0.5
0.4
0.3
0.2
0.1
0

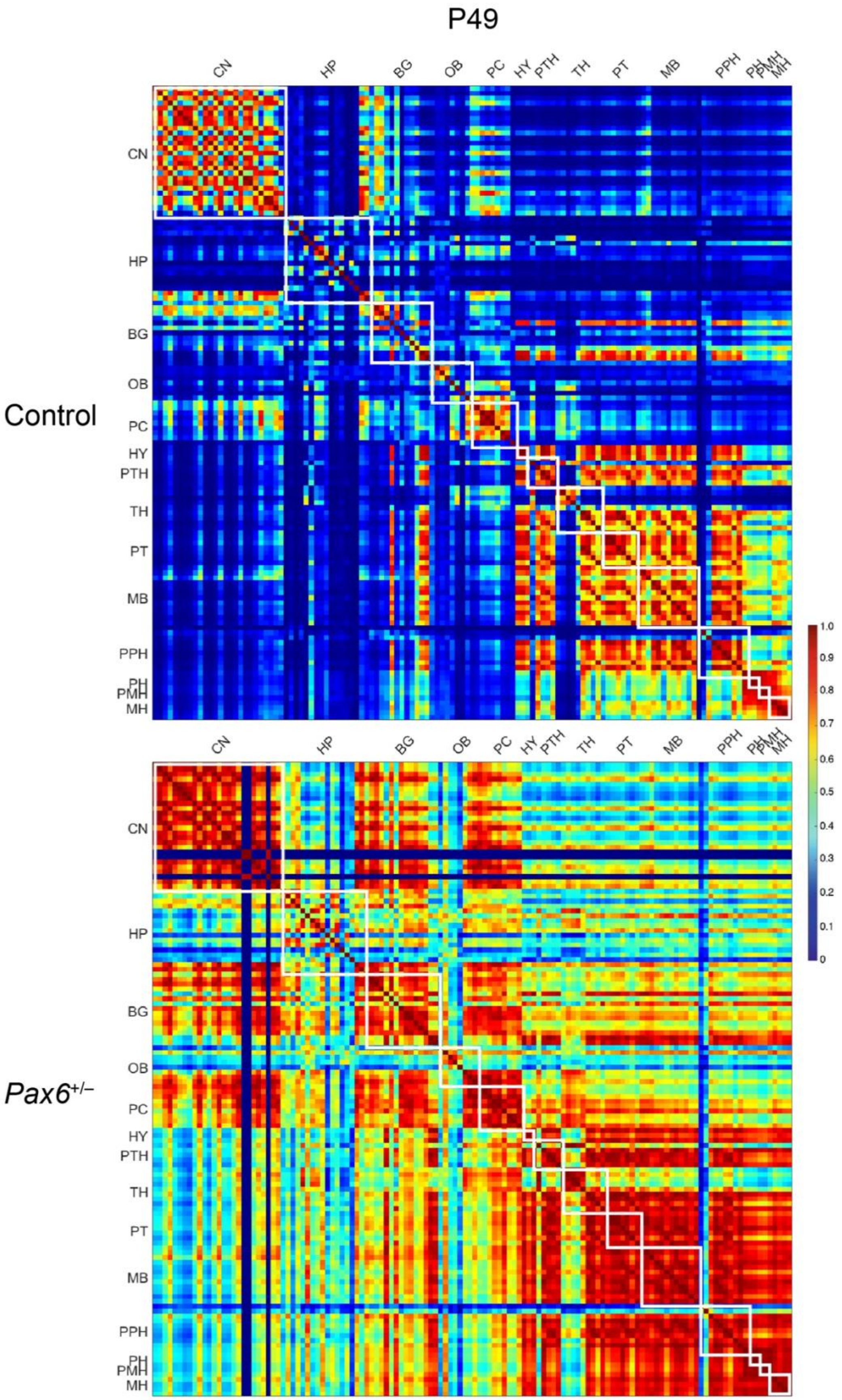

P49
Control
Pax6+/–
CN
HP
BG
OB
PC
HY
PTH
TH
PT
MB
PPH
PH
PMH
MH
1.0
0.9
0.8
0.7
0.6
0.5
0.4
0.3
0.2
0.1
0

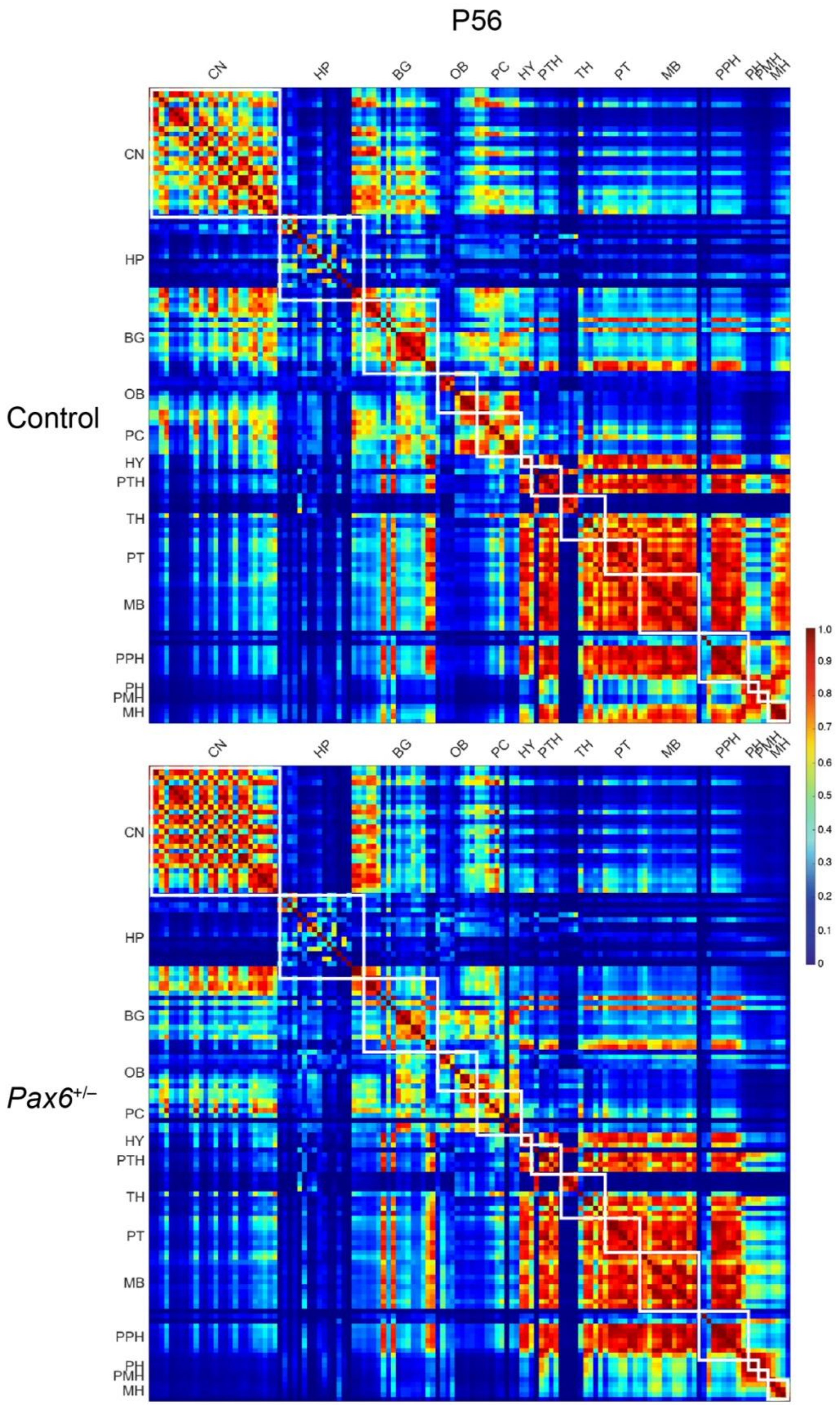
P56
Control
Pax6+/–
CN
HP
BG
OB
PC
HY
PTH
TH
PT
MB
PPH
PH
PMH
MH
1.0
0.9
0.8
0.7
0.6
0.5
0.4
0.3
0.2
0.1
0

**Figure S3. Brain subregion similarity matrices.**

Matrix heatmaps of similarities between pairs of subregions (rows and columns) at P1, P7, P14, P21, P28, P35, P42, P49 and P56 of control (top panels) and *Pax6*$^{+/-}$ (bottom panels) mice. White boxes indicate the subregions that belong to the same main brain region (see region list in Table S1). Scale bar, similarity values.

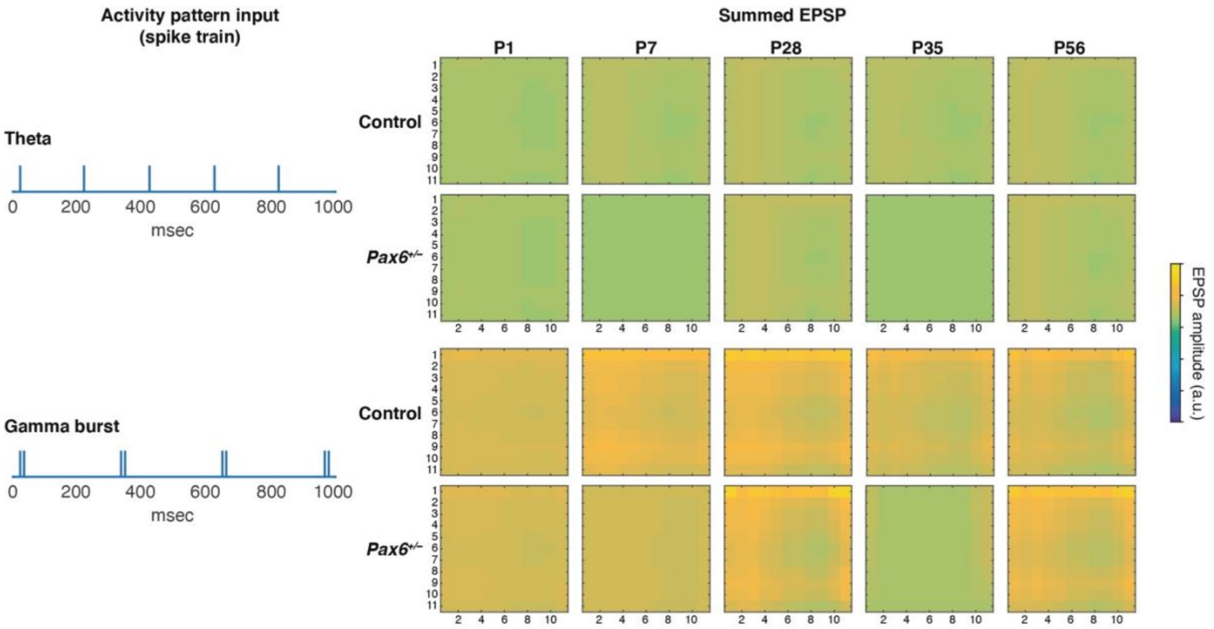


**Figure S4. Synaptic responses to physiological spike patterns**

Synaptic amplitudes of age groups (P1-P56) and genotype (control, *Pax6*$^{+/-}$) were scaled based on intensity (Table S14, S15). Synapses were activated by spike patterns representing theta (top) and gamma burst (bottom) activity. Summed EPSP response amplitudes were quantified (color bar, arbitrary units) and statistical differences between synaptic responses of control (upper) and *Pax6*$^{+/-}$ (lower) mice were assessed.

### Supplementary Table Legends

Table S1. Names of brain regions and subregions in heatmaps.

The identity of 131 subregions and their order used in heatmaps and similarity matrices.

Table S2. PSD95 synaptome parameters in different brain subregions.

Column A represents the 131 brain subregions analyzed and the first row the mouse ID number. PSD95 punctum density, intensity and size values in different brain subregions as defined in the Developing Allen Reference Atlas. Each spreadsheet contains data for each PSD95 parameter at one time point (P1, P7, P14, P21, P28, P35, P42, P49, P56). Abbreviations, see Table S1.

Table S3. SAP102 synaptome parameters in different brain subregions.

Column A represents the 131 brain subregions analyzed and the first row the mouse ID number. SAP102 punctum density, intensity and size values in different brain subregions as defined in the Developing Allen Reference Atlas. Each spreadsheet contains data for each SAP102 parameter at one time point (P1, P7, P14, P21, P28, P35, P42, P49, P56). Abbreviations, see Table S1.

Table S4. Colocalization of synaptome parameters in different brain subregions.

Column A represents the 131 brain subregions analyzed and the first row the mouse ID number. Colocalization values in different brain subregions as defined in the Developing Allen Reference Atlas. Each spreadsheet contains colocalization data at one time point (P1, P7, P14, P21, P28, P35, P42, P49, P56). Abbreviations, see Table S1.

Table S5. Subtype classification in different brain subregions at P1.

Column A represents the 131 brain subregions, the first row corresponds to the mouse ID number and the third row the number of the subtype. Subtype classification values in different brain subregions as defined in the Developing Allen Reference Atlas at P1. Each spreadsheet contains subtype classification data from each subtype (1-37). Abbreviations, see Table S1.

Table S6. Subtype classification in different brain subregions at P7.

Column A represents the 131 brain subregions, the first row corresponds to the mouse ID number and the third row the number of the subtype. Subtype classification values in different brain subregions as defined in the Developing Allen Reference Atlas at P7. Each spreadsheet contains subtype classification data from each subtype (1-37). Abbreviations, see Table S1.

Table S7. Subtype classification in different brain subregions at P14.

Column A represents the 131 brain subregions, the first row corresponds to the mouse ID number and the third row the number of the subtype. Subtype classification values in different brain subregions as defined in the Developing Allen Reference Atlas at P14. Each spreadsheet contains subtype classification data from each subtype (1-37). Abbreviations, see Table S1.

Table S8. Subtype classification in different brain subregions at P21.

Column A represents the 131 brain subregions, the first row corresponds to the mouse ID number and the third row the number of the subtype. Subtype classification values in different brain subregions as defined in the Developing Allen Reference Atlas at P21. Each spreadsheet contains subtype classification data from each subtype (1-37). Abbreviations, see Table S1.

Table S9. Subtype classification in different brain subregions at P28.

Column A represents the 131 brain subregions, the first row corresponds to the mouse ID number and the third row the number of the subtype. Subtype classification values in different brain subregions as defined in the Developing Allen Reference Atlas at P28. Each spreadsheet contains subtype classification data from each subtype (1-37). Abbreviations, see Table S1.

Table S10. Subtype classification in different brain subregions at P35.

Column A represents the 131 brain subregions, the first row corresponds to the mouse ID number and the third row the number of the subtype. Subtype classification values in different brain subregions as defined in the Developing Allen Reference Atlas at P35. Each spreadsheet contains subtype classification data from each subtype (1-37). Abbreviations, see Table S1.

Table S11. Subtype classification in different brain subregions at P42.

Column A represents the 131 brain subregions, the first row corresponds to the mouse ID number and the third row the number of the subtype. Subtype classification values in different brain subregions as defined in the Developing Allen Reference Atlas at P42.

Each spreadsheet contains subtype classification data from each subtype (1-37). Abbreviations, see Table S1.

Table S12. Subtype classification in different brain subregions at P49.

Column A represents the 131 brain subregions, the first row corresponds to the mouse ID number and the third row the number of the subtype. Subtype classification values in different brain subregions as defined in the Developing Allen Reference Atlas at P49. Each spreadsheet contains subtype classification data from each subtype (1-37). Abbreviations, see Table S1.

Table S13. Subtype classification in different brain subregions at P56.

Column A represents the 131 brain subregions, the first row corresponds to the mouse ID number and the third row the number of the subtype. Subtype classification values in different brain subregions as defined in the Developing Allen Reference Atlas at P56. Each spreadsheet contains subtype classification data from each subtype (1-37). Abbreviations, see Table S1.

Table S14. PSD95 gradients in the CA1sr of the hippocampus.

The PSD95 density, intensity and size parameters along the radial and tangential axis in the CA1 striatum radiatum subregion of the hippocampus. Each column contains the PSD95 values of individual mice. Each spreadsheet contains data for each PSD95 parameter at one time point (P1, P7, P14, P21, P28, P35, P42, P49, P56).

Table S15. SAP102 gradients in the CA1sr of the hippocampus.

The SAP102 density, intensity and size parameters along the radial and tangential axis in the CA1 striatum radiatum subregion of the hippocampus. Each column contains the SAP102 values of individual mice. Each spreadsheet contains data for each PSD95 parameter at one time point (P1, P7, P14, P21, P28, P35, P42, P49, P56).
